# Potential impacts of supplementing next generation long-lasting insecticidal nets with household-scale micro-mosaic deployment of indoor residual spraying with insecticides upon rates of incipient resistance trait emergence and selection

**DOI:** 10.64898/2026.08.18.745509

**Authors:** Dingani Chinula, Nicholaus Mziray, Neil Philip Hobbs, Busiku Hamainza, Thomas Reed, Samson Kiware, Gerry F. Killeen

**Affiliations:** National Malaria Elimination Centre, Chainama Hills Hospital Grounds, PO Box 32509, Lusaka, Republic of Zambia; School of Biological, Earth and Environmental Sciences, University College Cork, Cork, Republic of Ireland; Department of Environmental Health and Ecological Sciences, Ifakara Health Institute, Ifakara, Morogoro, United Republic of Tanzania; Department of Vector Biology, Liverpool School of Tropical Medicine, Liverpool, United Kingdom; Sustainability Institute, University College Cork, Cork, Republic of Ireland

## Abstract

Prolonged use of the few insecticide classes available for long-lasting insecticidal nets (LLINs) and indoor residual spraying (IRS) has driven widespread physiological resistance of malaria vector mosquitoes to this limited arsenal of active ingredients. However, recent innovations like next-generation LLINs (NG-LLINs) containing two complementary insecticides and new insecticide classes for IRS offer new opportunities for pre-emptive resistance management by deploying more diversified actives as mixtures, combinations, rotations or mosaics. Here a deterministic model of mosquito foraging behaviour was formulated to predict the probabilities of deterrence, mortality or successful feeding across repeated feeding attempts in scenarios with different combinations of NG-LLINs and/or IRS micro-mosaics with varying levels of insecticide diversification between neighbouring houses. Final fates were classified based on whether or not the mosquito eventually died or successfully fed, and whether the latter occurred indoors or outdoors after exposure to zero, one or several IRS insecticides. The primary outcome was the probability that a single *F₀* mosquito carrying a novel resistance trait to a new IRS insecticide successfully feeds, survives and reproduces, thereby establishing those traits within the population. The secondary outcome was the selection coefficient governing the spread of such novel resistance traits from the F₁ generation onwards. For highly anthropophagic and endophagic vectors like *Anopheles funestus*, combining NG-LLINs with IRS micro-mosaics using two insecticides may reduce emergence rates for novel resistance traits against IRS insecticides by approximately 2 to 2.5-fold, mainly through direct killing by NG-LLINs, although exposure to both IRS actives when forced to visit multiple houses also contributes to a lesser extent. However, such resistance management benefits are fundamentally constrained by outdoor feeding behaviours that limit or completely prevent indoor insecticide exposure. Increasing IRS micro-mosaic insecticide diversity beyond two actives is unlikely to further dampen resistance emergence rates because few mosquitoes survive long enough without feeding to encounter several IRS treatments. Once a resistance trait becomes established in the vector population, selection coefficients remain consistently high enough to force the spread of those traits, regardless of intervention combination. For more exophagic, zoophagic vectors like *An. arabiensis*, NG-LLINs plus IRS micro-mosaics are not expected to provide any meaningful resistance management benefit because frequent outdoor feeding, often on animals, allows them to largely avoid insecticide exposure altogether. Exclusively indoor-focused vector control strategies may not satisfactorily slow insecticide resistance emergence and spread, so new outdoor protection measures that close these coverage gaps with complementary insecticides will be needed.

## Introduction

Globally, the primary interventions for malaria vector control are long lasting insecticidal nets (LLINS) and indoor residual spraying (IRS) of long-lasting insecticide formulations [1, 2]. For many years, only four chemical classes of insecticide (pyrethroids, organochlorines, carbamates and organophosphates) have been available for vector control, with pyrethroids being the only class approved for LLINs until relatively recently [1, 2]. The prolonged use of these few insecticide classes exerted considerable selection pressure on vector populations, contributing to the emergence and widespread distribution of physiological resistance across sub-Saharan Africa [3-5]. In response to this, WHO published the Global Plan for Insecticide Resistance Management (GPIRM)[1]. The GPIRM suggested that the future use of rotations, mosaics, combinations and mixtures as viable insecticide resistance management (IRM) strategies once new actives became available, primarily informed by broad evolutionary principles and process-explicit mathematical modelling rather than empirical evidence.

Similarly to many other African countries, Zambia considers LLINs to be the standard minimum best practice for vector control, which is selectively supplemented with IRS in areas with high transmission intensity [6]. To ensure sustained coverage, mass LLINs distribution are implemented every three years across Zambia and began in 2005 [7]. To address the growing threat of insecticide resistance, Zambia first switched to LLINs co-treated with both a pyrethroid and the pyrethroid synergist piperonyl butoxide (PBO) since 2020 [8] and will now switch to NG-LLINs in the mid-2026 [9]. However, each switch to a new LLIN type has been implemented on a reactionary basis, with pyrethroids remain as one of the active ingredients regardless of the LLIN product deployed. Correspondingly, pre-existing pyrethroid resistance could potentially undermine their immediate vector control and longer-term insecticide resistance management (IRM) functions [10].

Zambia has also deployed IRS at scale with considerable success [11]. Between 2000 and 2012, Zambia implemented IRS with pyrethroids, carbamates and organo-chlorines [11, 12]. However, due to insecticide resistance [11-15] and limited alternatives, the country switched to an organophosphate from 2013 to 2017 [11, 16]. From 2018 up to 2025, the neonicotinoid clothianidin and a co-formulation of clothianidin with deltamethrin have been deployed as the default products for IRS, although DDT was also used focally between 2020 and 2022 before being completely withdrawn due to environmental concerns [8, 9]. This continuing pattern of excessive, extended reliance on a small handful of insecticide classes, coupled with a reactionary IRM approach may well result in widespread resistance to all of them within a few short years unless more effective IRM practices are promptly put in place.

In years since GPIRM was published, the availability of new public health insecticides classes (e.g. pyrroles, meta-diamides and neonicotinoids) and formulations (e.g. mixtures) offers greater chemical diversity for combatting insecticide resistance [17]. This presents an opportunity for malaria control programmes to start adopting pre-emptive resistance management strategies and tactics [1, 4, 18, 19] if they are bold enough to seize it promptly. IRM is most effective when implemented early, before resistance becomes prevalent or, more ideally, before it even emerges [1, 3, 4, 18, 19]. This is particularly important because there is no guarantee that new active ingredients will continue to become available quickly enough to offset the rate at which existing actives are lost to resistance. It is therefore crucial that any such new actives are astutely deployed in a forward-looking manner [4, 18, 20, 21], to avoid past mistakes that compromised long-term sustainability of vector control interventions [4, 18, 21]. The urgent need to embrace pre-emptive resistance management strategies, to preserve the efficacy of existing vector control measures [4, 18, 21], is therefore reinforced by the fleeting current opportunity to do so: It is indeed most probably a case of now or never.

GPIRM makes no reference to spatial scales when referring to *mosaics*, making no distinction between practices in which the insecticide classes used are varied between neighbouring villages (Figure 1A), or even at coarser district or provincial scales, and micro-heterogeneous deployment as household-scale *micro-mosaics* with different insecticides applied to neighbouring houses within the same village (Figure 1B). These differences in the scale of deployment heterogeneity may impact IRM effectiveness because mosquitoes may be exposed to several IRS active within a single village in the latter scenario (Figure 2) but not the former. By contrast, mathematical models suggest that rotations involving swapping of active ingredients each year and household-scale micro-mosaics (Figure 1B) may have comparably modest effects upon emergence and fixation timelines [22].

**Figure 1.**
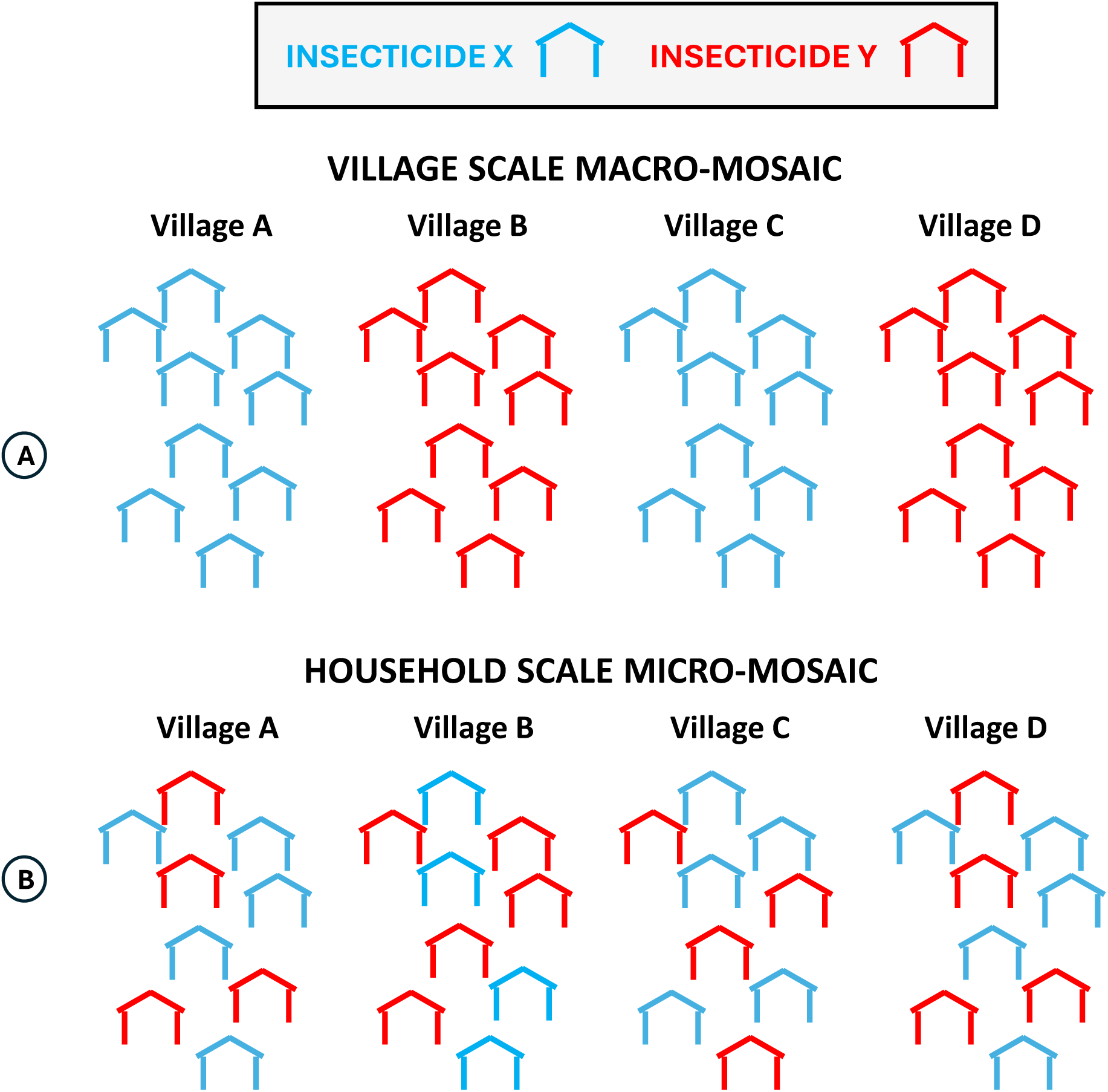
A schematic illustration of (A) macro-mosaic IRS application of different insecticides in different villages versus (B) micro-mosaic application at with different insecticides in different neighbouring houses within individual villages [23].

**Figure 2.**
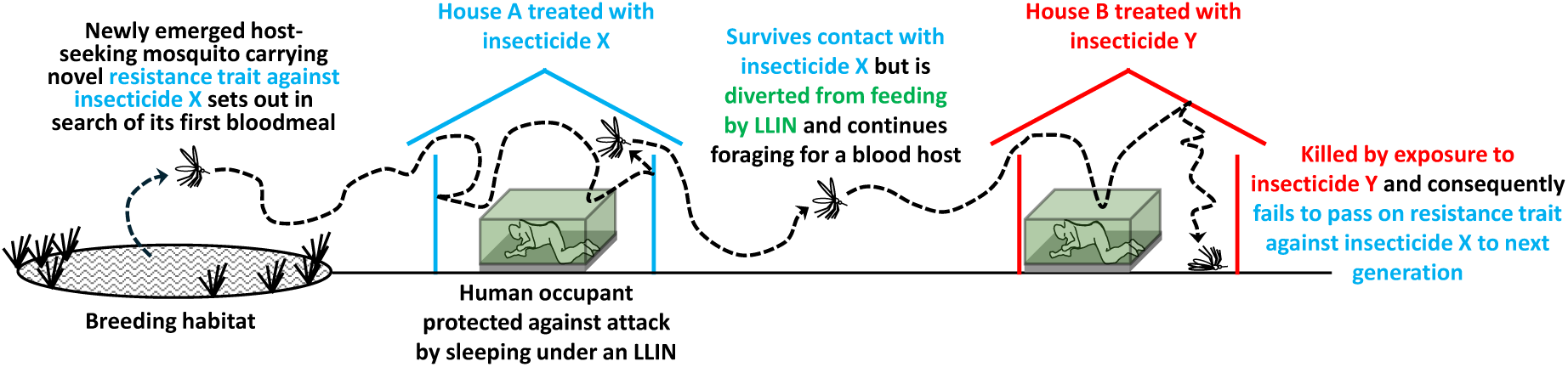
A schematic illustration of how mosquitoes may be exposed to different insecticide formulations in different houses within a single gonotrophic cycle in a village where NG-LLINs are supplemented with household-scale micro-mosaic IRS deployment of two distinct and complementary insecticides [23] as per figure 1B.

However, these particular models assumed that mosquito vectors resting inside human habitations do so only once during each fully concordant gonotrophic cycle, thereby encountering only a single insecticide each time it feeds, gestates and then lays eggs [22, 24]. Given the currently high coverage of LLINs in Zambia and elsewhere across much of Africa, which frequently force mosquitoes to move from one house to the next in search of a blood meal, this assumption may underestimate the potential impact of micro-mosaics in relevant contemporary contexts [2, 25].

Mosquitoes entering human dwellings where the occupants use LLINs are frequently killed or deterred from feeding, so those that survive are often forced to visit multiple households within a single gonotrophic cycle in order to feed and then reproduce successfully (Fig 2) [26-28]. The evasive behaviours expressed by mosquitoes in response to high coverage with LLINs substantially alter their life histories, increasing the likelihood of encountering two or more of the different insecticides used in micro-mosaic deployment formats for IRS (Figure 2). Correspondingly, such extended foraging bouts may reduce the probability that any mosquito born with a heritable resistant trait against any single active ingredient might pass on the causative allele(s) to subsequent generations.

The study therefore explores how NG-LLINs combined with micro-mosaic IRS may be used for IRM by ensuring the parental generation (*F_o_*) carrying any novel heritable alleles are exposed to at least two different active ingredients from different classes (Figure 2). The overall goal of this IRM tactic is to reduce the probability that *F_0_* mosquitoes may pass an incipient resistance trait on to the next (*F_1_*) generation than would otherwise be the case, rather than try to slow the selection of such traits to high frequency once their progeny are established in the population in subsequent generations.

## Methods

### Conceptual framework and outcomes of interest

As micro-mosaics are still in the early stages of evaluation, either empirically or theoretically, there remains much uncertainty regarding how to realistically model their potential role in IRM. As far as we are aware, there have been only two prior modelling frameworks that explicitly model micro-mosaic for mosquito control [22, 24, 29], working on the hypothesis that micro-mosaics may add a “temporal-mixture” effect as foraging female mosquitoes encounter different insecticides across multiple contacts within each gonotrophic cycle. These models focused on the complexity in the genetics (monogenic, polygenic) and selection dimension (including dominance, insecticide decay, cross resistance) which necessitated reductions in behavioural and/or mosquito dynamics complexity, which limited the diversity of insecticides (up to two) which could be simultaneously “deployed” in the model (at any time point) as either LLINs or IRS. In this new modelling framework for micro-mosaics, we instead focus on the behavioural complexity and the number of insecticides potentially simultaneously deployed, but this necessitates reducing the genetic complexity.

More importantly, however, this model explores the varying consequences of different specific vector control intervention tactics in the context of a truly pre-emptive IRM strategy, meaning that all the insecticides being used are entirely new and no pre-existing resistance traits are already established within the vector population [19]. This theoretical exercise consequently focuses on the prevention of incipient resistance traits from establishing themselves in the population in the first place, by killing the first single individual *F_0_* mosquito carrying a novel heritable resistance trait before it can reproduce. The selection pressures that favour subsequent spread of that resistant trait throughout the population from *F_1_* onwards are also considered, but our primary focus is on the *F_0_* generation. This distinction between the early establishment versus later spreading phase is conceptually analogous to wildfire dynamics. Initial sparks may either dissipate on contact with the ground or perhaps ignite a small fire that may go unnoticed for quite some time. But once established, a small fire can quickly grow into a ragging infernal that is patently obvious but vastly more difficult to control (Box 1).

#### Box 1. Conceptualizing trajectories of insecticide resistance emergence and evolution in terms of an analogy with sparks and subsequent wildfires in a forest, all of which can be related to the core underlying assumptions of the model (Box 2, Figure 3).

An analogy may be made between insecticide resistance evolution and wildfires, which may begin with unseen tiny *sparks* (Yellow initial seed events in figure 3) that can potentially ignite a self-sustaining *fire* that subsequently becomes obvious only after it spreads (Orange and then red trajectories in figure 3). Although this modelling analysis does consider the latter selection pressures that fan the evolutionary flames of such established evolutionary wildfires towards high frequency across entire populations in later *F_≥1_* generations (Orange trajectories that progress to red ones in figure 3), the primary focus of this exercise is former sparks of initial emergence events at the scale of single individual *F_0_* mosquitoes carrying the very first and only original copy of the relevant resistance alleles that start those wildfires in the first place (Yellow initial seed events in figure 3).

The primary outcome considered in this theoretical study is therefore the estimated probability that such an individual mosquito carrying a heritable novel resistance trait against a single new IRS insecticide will survive and reproduce, accounting for fates that involve exposure to no new IRS insecticides or only one (*P_ϕ,≤1_*), reflecting the rate at which sparks occur among *F_0_* generation mosquitoes and subsequently lead to the establishment of a derived *F_1_* generation that represents the established beginnings of a wildfire. The secondary outcome considered herein is the estimated value for the selection coefficient (*s*), a widely used and robustly standardized evolutionary biology parameter that reflects the evolutionary pressures favouring selection of that resistance trait once established in the population from *F_1_* onwards, analogous to the rate at which a wildfire grows exponentially until it consumes the entire forest.

Crucially, this model is based on four core assumptions about the biological and intervention delivery circumstances under which IRS is deployed in combination with NG-LLINs (Box 2), all of which may be related to our analogy with how wildfires begin and then progress (Box 1).

#### Box 2. The four core assumptions made for this modelling exercise regarding what a pre-emptive insecticide resistance management strategy actually means in biological terms.

1. In the absence of any positive selection pressure from insecticide use to maintain them in the population over the long term, sporadically occurring resistance traits die out entirely through negative selection and genetic drift within a few short generations. In such a hypothetical baseline scenario, preceding the introduction of these IRS and NG-LLINs with combinations of entirely new active ingredients on a truly pre-emptive basis, any such sporadically occurring emergent resistance traits are not subsequently maintained as a stable component of heritable standing variation at any non-zero frequency whatsoever within the mosquito population (Figure 3, top panel). This assumption is justified on the basis that new insecticides, specifically those developed for genuinely pre-emptive use in the strictest sense, will necessarily have novel modes of action against which no pre-existing resistance traits occurs among wild populations.
2. Before and after implementation of a truly pre-emptive IRM intervention strategy, resistance traits against these new insecticides arise spontaneously in individual *F_0_* mosquitoes only very occasionally, meaning once every few years rather than within days, weeks or months. This assumption is justified on the basis of prior historical experience with a range of existing IRS and LLIN insecticides that have been successfully used to entirely eliminate highly anthropophagic and endophagic vectors from a diversity of settings across the tropics [30], noting that failures to eliminate target mosquito species usually arise from behaviours of mosquitoes and people that undermine the crucial mass populations effects of these key vector control measures [30-33]. None of these local extinction events, all of which took at least two years before the vector in question finally disappeared [30], could have occurred if robust resistance traits pre-existed within the standing variation of those populations or spontaneously arose de novo within the period taken for that mosquito population to entirely fizzle out. Similarly, experience with some antimalarial drugs is encouraging: Sheer luck and very low rates of relevant mutations allowed extended use of chloroquine monotherapy for decades [34], while pre-emptive deployment of artemisinin combination therapies [35-37] have yielded effective product lifetimes in excess of 20 years although emerging resistance now necessitates reactive tactics [38-40]. This assumption is also justified on the basis of the likely futility of almost any IRM strategy otherwise: Unless genuinely pre-emptive IRM strategies can extend insecticide product lifetimes by years or decades, they will simply not be useful enough to be worthwhile.
3. All active ingredients used for these combinations of IRS with NG-LLINs are introduced in a truly pre-emptive manner, meaning before any resistance traits to the relevant insecticides have been allowed to emerge and be selected for by previous use as stand-alone active ingredients or supplementary active ingredients to those to which resistance is already widespread [19]. The hypothetical scenarios explored herein therefore reflect truly pre-emptive strategies that have yet to be embraced at policy level or implemented at scale in practice [19]. The *a priori* hypothesis underpinning this assumed policy and practice scenario is that such pre-emptive strategies will be required to maximize the useful market lives of new insecticides in the future, to not only make optimal use of new products being developed for the malaria vector control market for as long as possible (Figure 3, bottom panel versus middle) but also incentivize such investments in innovation by minimizing risk and maximizing financial viability for manufacturers over the longer term.
4. Novel resistance traits first arise in a single individual *F_0_* mosquito borne into a brood of otherwise fully susceptible siblings with a fully susceptible mother and father, so any vector control intervention that substantially reduces the probability of such a single resistant *F_0_* mosquito surviving and reproducing will often decisively terminate that resistant lineage at its most vulnerable stage, namely that *F_0_* generation when it consists only of a single insect (Figure 3, bottom panel), rather than the larger numbers that may occur in subsequent *F_≥1_* generations that have much higher chances of yielding at least one offspring that survives and reproduces despite those increased adult mortality risks. Crucially, the *F_0_*, *F_1_* and *F_>1_* referred to throughout this manuscript respectively represent the first single insect carrying a particular incipient resistant trait, first subsequent generation of that lineage and then all derived generations of progeny, rather than the first generation that has been exposed to a pre-emptive IRM intervention *per se*, regardless of heritable resistance status.

Regarding the details of how the consequences of one intervention scenario differ from another, this deterministic model calculates the probabilities of sequential outcomes of mosquito interactions with vector control insecticides, deployed as a combination of NG-LLINs and household scale micro-mosaics of IRS with two (Figure 1B) or more complementary insecticide formulations. This model particularly focuses on the how the varying probabilities of deterrence, mortality and successful feeding fates are influenced by the use of NG-LLINs, and how this affects the consequent likelihood of mosquitoes being exposed to one or more IRS insecticides over the course of a successfully completed first gonotrophic feeding cycle (Figure 4).

**Figure 3.**
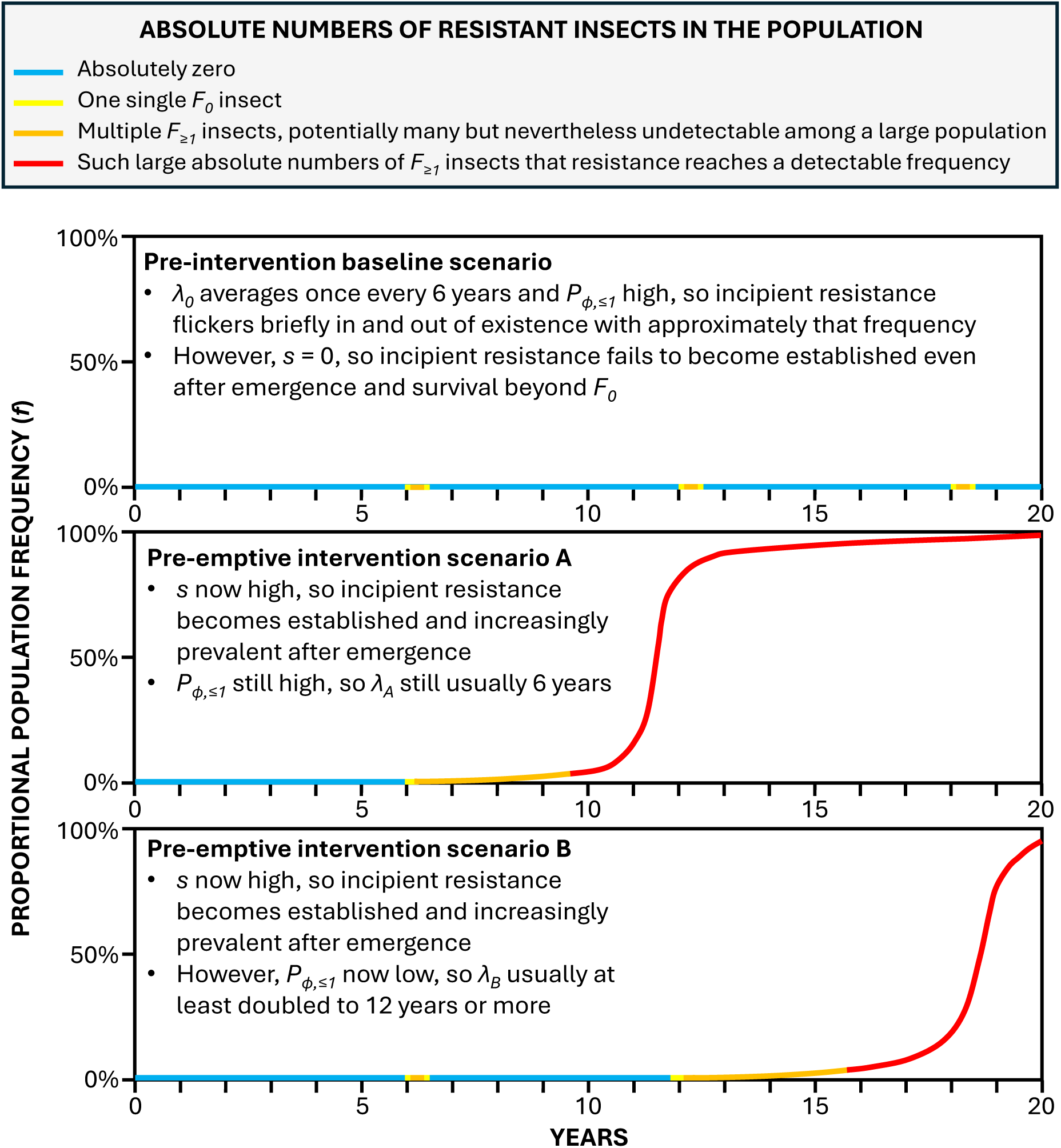
A schematic illustration of the core assumptions made regarding the population dynamics of the mosquito population under a pre-intervention baseline scenario (Top panel) and two contrasting pre-emptive insecticide resistance management intervention scenarios that do (Bottom panel) and do not (Middle panel) substantially reduce the survival and reproduction probability of individual resistance insects. See boxes 1 and 2 for further explanation.

**Figure 4.**
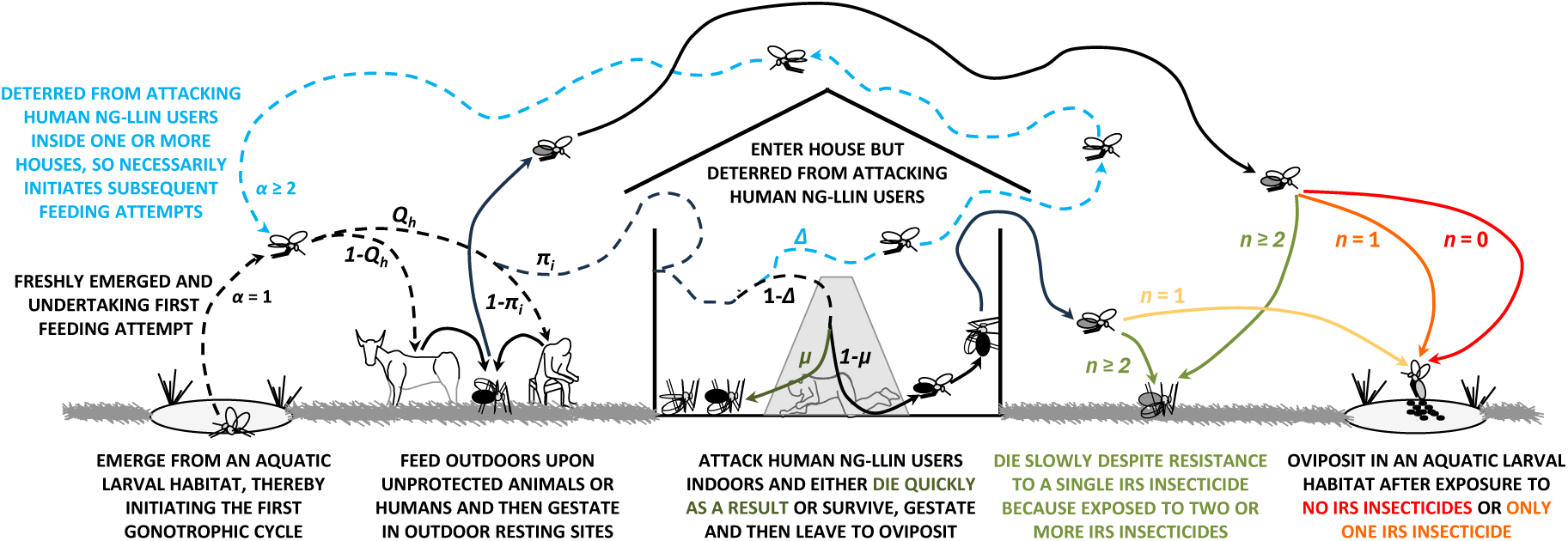
An explicit schematic representation of the mathematical model presented herein for all the possible foraging, blood acquisition, gestation and oviposition behaviours exhibited by malaria vector mosquitoes over the course of its first gonotrophic cycle, which may require the mosquito to make two or more feeding attempts before it may reproduce successfully. As annotated in this schematic, distinct possible fates arising from any given feeding attempt are classified as either deterrence and diversion (*Δ*), mortality (*μ*) or successful feeding (*ϕ*). See Table S1, Supplementary File 1 for fully detailed definitions and explanations of the symbols used. Note that the colour scheme used here, in subsequent figures, and in Table S2, Supplementary File 1 is unrelated to that used in figures 1 to 3.

The primary outcome is therefore the proportion of mosquitoes that successfully feed and survive any indoor encounters with NG-LLINs during their first gonotrophic cycle after having been exposed to one, and only one IRS insecticide (*P_ϕ,1_*) or none as defined in Table S1, Supplementary file 1. This primary outcome reflects the predicted probability that a mosquito born with an incipient resistant trait against that single IRS insecticide to which it was exposed will successfully survive and thus pass on its genes. The secondary outcome *s* is the fraction of all mosquitoes surviving to complete their first blood meal that have been exposed to only one IRS insecticide, rather than none (Table S1, Supplementary File 1). Whilst our model is not genetically explicit, this is effectively equivalent to a selection coefficient as per the established population genetics literature [41, 42], which can be defined as the proportional disadvantage of the less-fit type (here susceptible mosquitoes) relative to the fitter type (here mosquitoes resistant to that single class of IRS insecticide). Our equation for *s* incorporates all possible behavioural/exposure pathways (see Figure 4 and Tables S1 and S2 in Supplementary File 1) via which a putative resistant type can end up contributing more to the next generation than a putative susceptible type, thus favouring the propagation of the resistance trait (and the implicit underlying resistance allele(s)) in the population from the *F_1_* generation onwards. This secondary outcome is an indicator of the selection pressure favouring subsequent spread throughout the population of any resistance traits against individual insecticides that may occur. In terms of the analogy described in Box 1, the primary outcome describes the probability of sparks occurring, while the secondary outcome relates to the likelihood and intensity of any subsequent wildfire.

However, a number of other endpoints and outcomes are possible, especially when the possibility that NG-LLINs may force mosquitoes to undertake multiple feeding attempts (Figure 4) is considered. Correspondingly, a mosquito may experience a fate (*f*) of being deterred from feeding, being killed in the attempt, or eventually succeeding in feeding (*f* = *Δ*, *μ* or *ϕ*, respectively) in an indoor or outdoor location (*l* = *i* or *o*, respectively) after exposure to *n* out of all the *N* different insecticides deployed in an IRS micro-mosaic over up to four feeding attempts (α = 1, 2, 3 or 4). The probabilities of all these numerous possible intermediate and final fates may therefore be expressed as the generic term *P_f_*_,*l*,*n*,*α*_ as per Table S1 in Supplementary File 1. All the different possible outcomes of host seeking and blood acquisition attempts that often have to be repeated are most easily conceptualized by first considering the first feeding attempt (First column of equations in Table S2, Supplementary File 1). It is then relatively straightforward to consider the consequences of mosquitoes having to repeatedly gamble with the same opportunities and risks when their second, third and fourth attempts to feed fail on one or more occasions (Second, third and fourth columns of equations in Table 2 of Supplementary File 1, respectively).

Fully explicit details of the mathematical model developed and applies for this purpose are provided in Supplementary File 1. Similarly, the rationale and specific numerical values chosen for parameterizing the model based on reasonable stereotypes of two behaviourally distinct categories of mosquitoes within a typical African malaria vector guild are fully outlined in Supplementary File 1.

## Results

For highly anthropophagic and endophagic vectors like *An. funestus*, most mosquitoes (66%) successfully obtain a blood meal on their first feeding attempt in the absence of bed nets (Figure 5, panels A, E, I, M). In contrast, untreated bednets, NG-LLINs with excito-repellent properties and NG-LLINs with purely lethal modes of action are expected to reduce successful feeding indoors at the first attempt to 29%, 3%, and 6.5%, respectively, regardless of the levels of IRS insecticide diversity deployed (Figure 5, all panels other than A, E, I and M).

**Figure 5:**
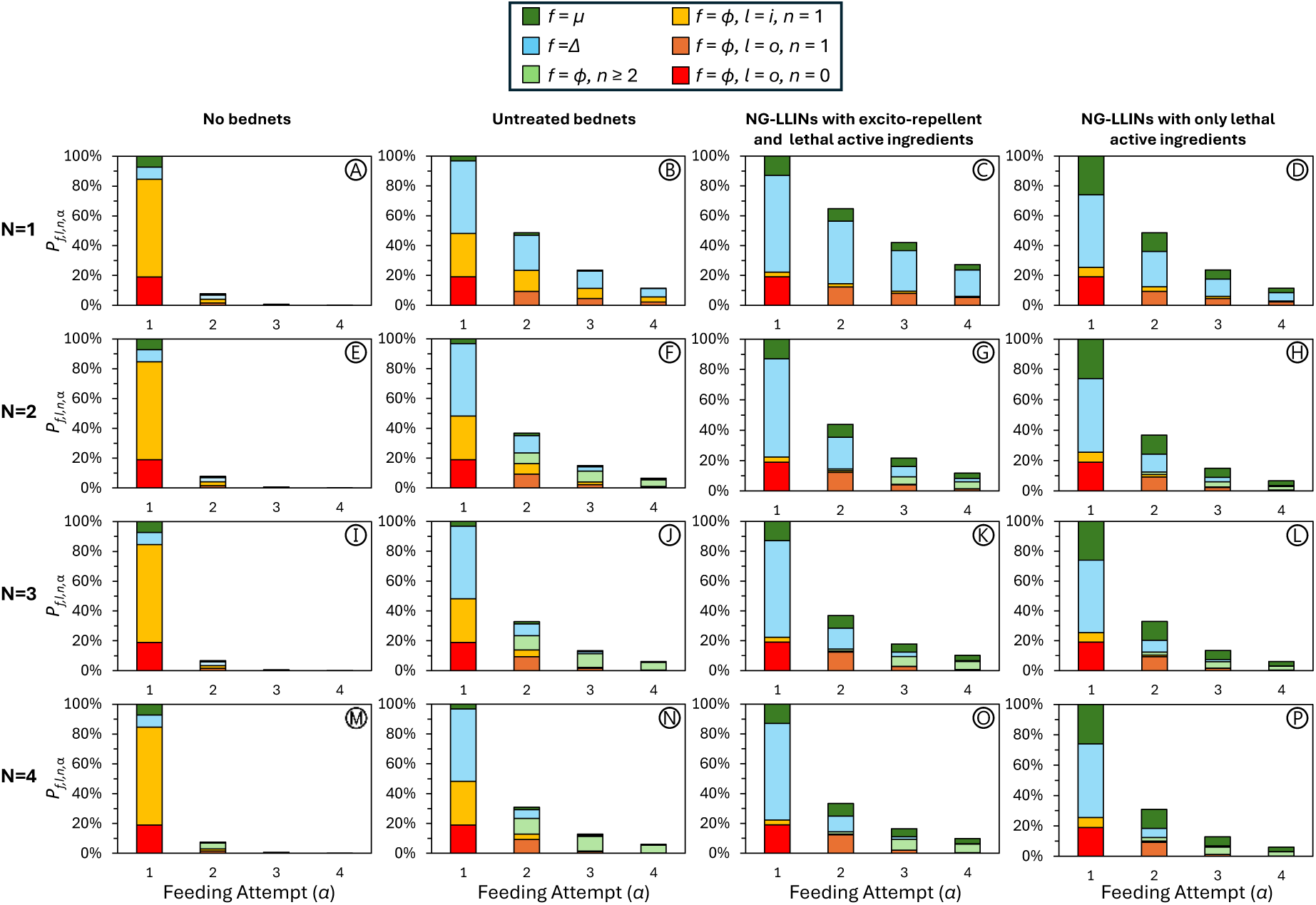
The expected proportions of host-seeking *An. funestus s.s.* mosquitoes starting a gonotrophic cycle (*P_f,l,n,α_*) that experience a deterred, died or successfully fed fate (*f = Δ*, *μ* or *ϕ*, respectively) at either an indoor or outdoor location (*l* = *i* or *o*, respectively) after exposure to none, one or several of the insecticides used for indoor residual spraying (IRS; *n* = 0, 1 or ≥2) in the first, second, third or fourth feeding attempt (*α* = 1, 2, 3 or 4) for all possible intervention scenarios combining either no bednets, untreated bednets, next generation long-lasting nets (NG-LLINs) exhibiting both lethal and excito-repellent properties or NG-LLINs exhibiting only lethal modes of action with IRS using either blanket coverage with only one insecticide (*N* = 1) or household-scale micro-mosaic formats with two to four insecticides (*N* = 2, 3 or 4).

For NG-LLINs with excito-repellent properties, the dominant effect is strong behavioural deterrence, with approximately 65% of mosquitoes repelled during the first feeding attempt (Figure 5, panels C, G, K, and O). Even among those expected to subsequently feed successfully on the second attempt, the vast majority subsequently feed outdoors after encountering a single IRS insecticide indoors. These mosquitoes are particularly relevant to IRM, as they are likely contributors to the propagation of any incipient resistance traits beyond *F_0_* due to their incomplete exposure to any of the diversity of IRS actives deployed as micro-mosaics.

By contrast, NG-LLINs with purely lethal properties induce substantial mortality, which accumulates across successive feeding attempts, totalling 49% overall (Figure 6, panels A, C, E, and G). This cumulative mortality markedly reduces the pool of mosquitoes that could possibly pass on resistance traits into subsequent generations (Figure 5, panels D, H, L, and P). Such reductions reflect strong suppression of “spark generation,” whereby resistant *F₀* mosquitoes are killed before they can successfully blood-feed, survive, and reproduce (*P_ϕ,≤1_*). Crucially, however, even for such NG-LLINs lacking excito-repellent properties, most of the mosquitoes that are exposed to only one IRS active ingredient inside houses and then go on to feed successfully ultimately do so outdoors.

**Figure 6:**
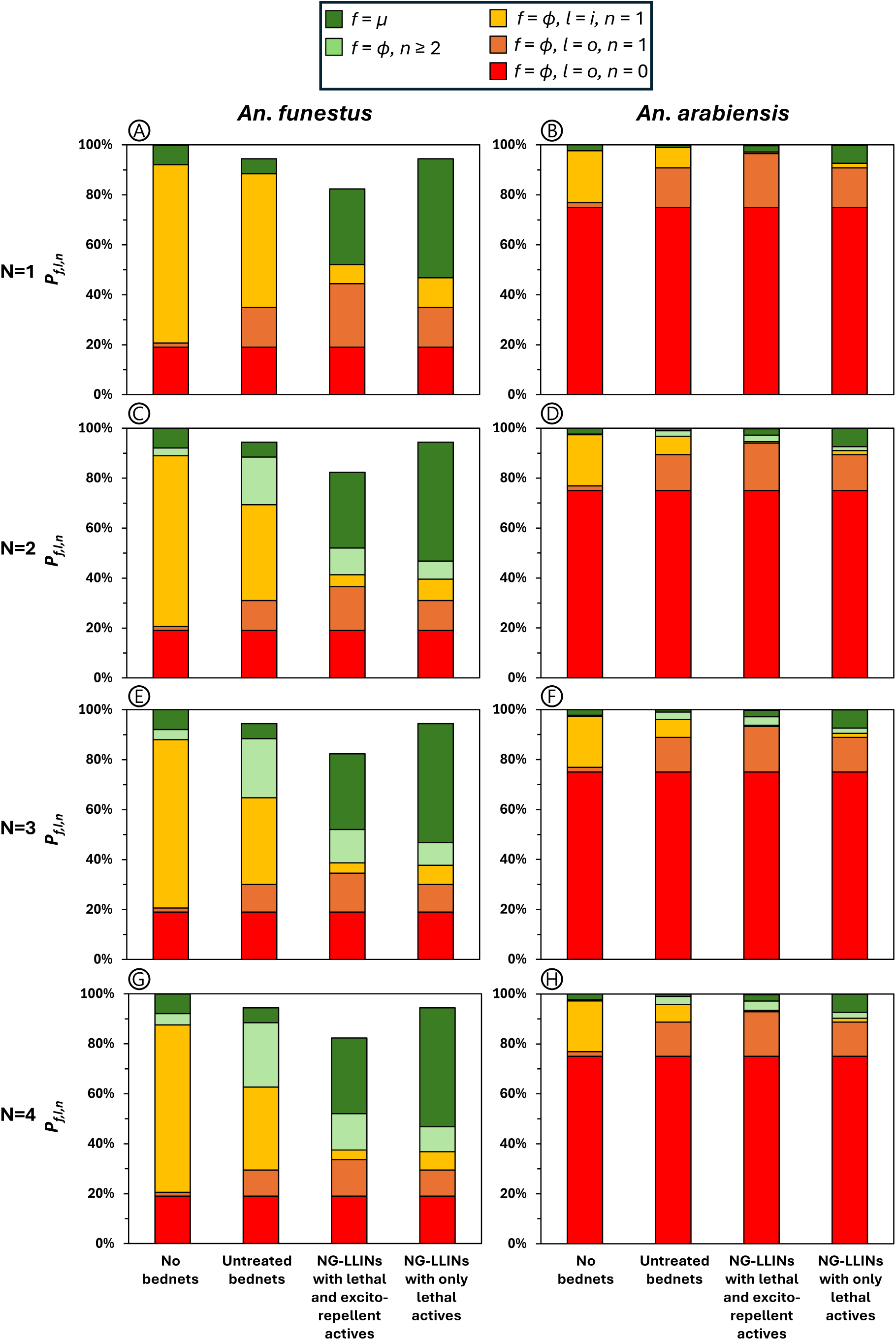
The expected cumulative proportions of host-seeking mosquitoes starting a gonotrophic cycle (*P_f,l,n,α_*) that experience a deterred, died or successfully fed final fate (*f* = *Δ*, *μ* or *ϕ*, respectively) at either an indoor or outdoor location (*l* = *i* or *o*, respectively) after exposure to none, one or several of the insecticides used for indoor residual spraying (IRS; *n* = 0, 1 or ≥2) over up to four feeding attempts (summed over α = 1, 2, 3 and 4) for all possible intervention scenarios combining either no bednets, untreated bednets, next generation long-lasting nets (NG-LLINs) exhibiting both lethal and excito-repellent properties or NG-LLINs exhibiting only lethal modes of action with IRS using either blanket coverage of only one insecticide (*N* = 1) or household-scale micro-mosaic formats with two to four insecticides (*N* = 2, 3 or 4).

Regardless of NG-LLIN properties, the vast majority of fates relevant to resistance emergence and evolution trajectories occur after only one or two feeding attempts. From the third to fourth feeding attempts, surviving fractions of unfed mosquitoes decline sharply and approach zero (Figure 5), so the possibility of surviving repeated failed feeding attempts is expected to be so rare that it is of negligible biological significance regarding IRM. The majority of all the possible fates that may determine the success or failure of these various approaches to IRM involve only one feeding attempt, with the almost of all of the remaining fates that matter involving only two feeding attempts (Figure 5).

This simple theoretical observation that most of the expected fates that matter occur after only one or two feeding attempts (Figure 5) has important practical implications for optimizing the degree of insecticide diversity used in IRS micro-mosaics. Specifically, it indicates that very few mosquitoes will survive enough feeding attempts inside houses to encounter more than two IRS insecticides in a given gonotrophic cycle, regardless of how many distinct IRS actives are deployed across the village they are foraging within (Figure 5). Mortality prior to successful feeding is the dominant expected effect of NG-LLINs in these simulations, regardless of whether or not they have excito-repellent properties, thus limiting the extent to which they may force mosquitoes into repeated encounters with multiple IRS insecticides (Figure 5). Consequently, increasing IRS insecticide diversity beyond two active ingredients in a given time and place yields diminishing returns (Figures 6 and 7) because so few mosquitoes are expected to survive long enough to experience sequential exposures to IRS-treated surfaces inside multiple houses (Figure 5).

**Figure 7:**
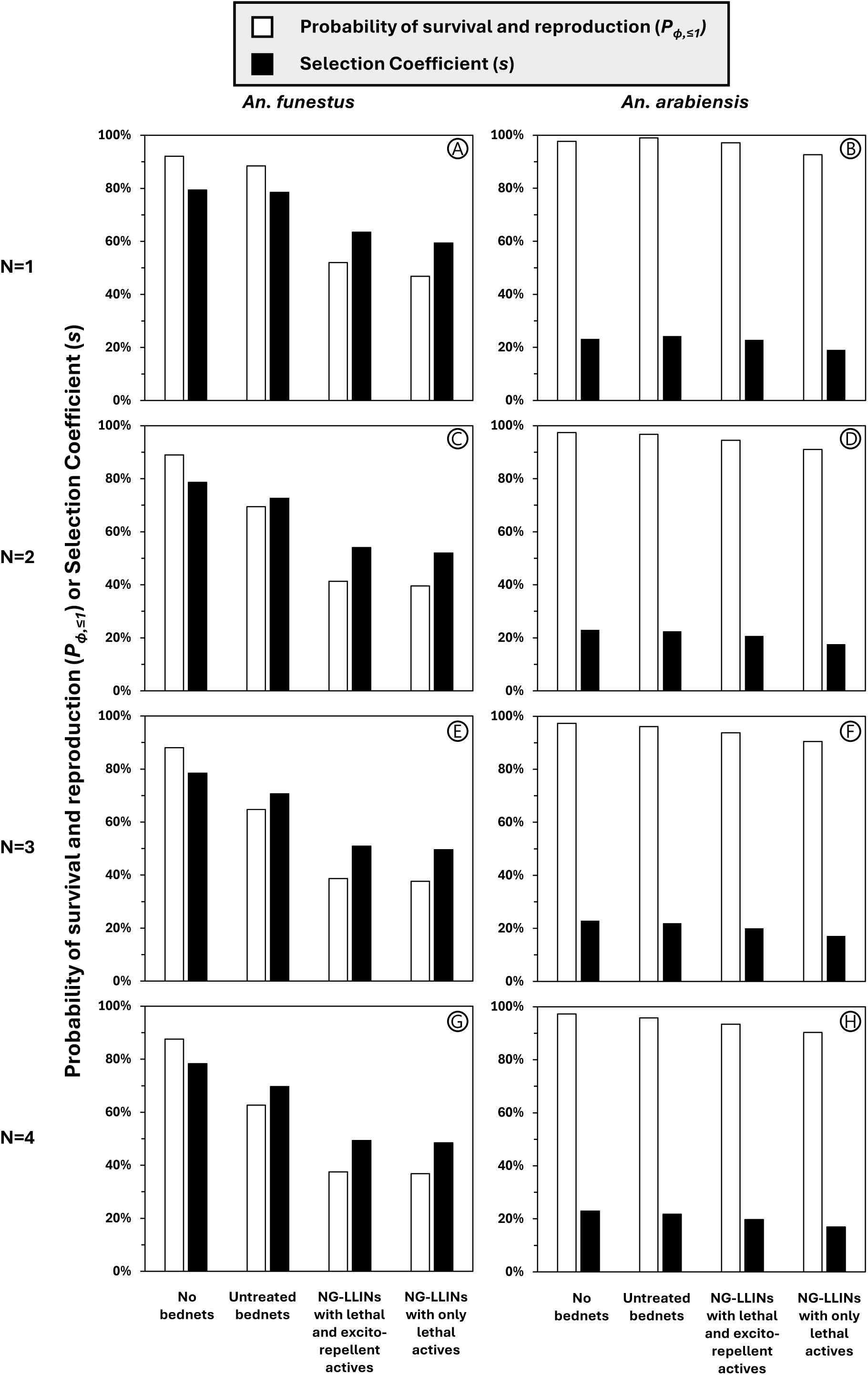
The calculated probabilities of feeding, survival and successful reproduction (P*_ϕ,≤1_*; Equation 7.3, Table S2, Supplementary File 1) and selection coefficients (*s*; Equation 8, Table S2, Supplementary File 1) for both *An.funestus s.s.* and *An. arabiensis* after exposure to all possible intervention scenarios combining either no bednets, untreated bednets, next generation long-lasting nets (NG-LLINs) exhibiting both lethal and excito-repellent properties or NG-LLINs exhibiting only lethal modes of action with IRS using either blanket coverage with only one insecticide (*N* = 1) or household-scale micro-mosaic formats with two to four insecticides (*N* = 2, 3 or 4). Using the wildfire analogy outlined in Box 2, the former primary outcomes is proportional to the rate at which the sparks of incipient resistance arising in individual *F_0_* generation mosquitoes yield larger numbers of progeny in subsequent *F_≥1_* generations. The latter secondary outcome represents the relative survival and reproduction advantage of those *F_≥1_* progeny that drives the steady selection of the resistance alleles they carry to high population frequency, reflecting the strength of the winds that fan the flames of such an established wildfire in the analogy outlined in Box 2.

The overall contributions of all the fates that arise from all simulated numbers of possible feeding attempts are summarized in figure 6 and presented in terms of the overall primary and secondary outcomes in figure 7. NG-LLINs and, to a lesser extent, supplementing them with IRS diversification at household level may indeed reduce the absolute probability of resistance emergence beyond *F_0_* at any given time (*P_ϕ,≤1_*), as well as the degree of selective pressure favouring any such traits that do establish themselves in subsequent *F_≥1_* generations (*s*) (Figure 7). However, they fall far short of eliminating such risks: Based on the expected modest reductions of *P_ϕ,≤1_* that are illustrated in figure 7, it is reasonable to expect that the emergence of such resistance traits may be delayed considerably but not that it is prevented indefinitely. Also, once resistance becomes established in the *F₁*and subsequent generations, the lower but nevertheless substantial simulated values for *s* imply that those traits may be expected to spread rapidly thereafter.

Expressing the *P_ϕ, ≤1_* estimates in figure 7 in relative reciprocal terms, expected values for the relative duration of the expected mean lifetime for pre-emptive use of an IRS insecticide for IRS micro-mosaics alone (*Λ_Ω,0_*, where *Ω* and *0* respectively represent scenarios where *N* ≥ 2 versus 1 in the absence of any bednets) all approximate to 1.04, suggesting little impact on resistance evolution trajectories in practical terms, regardless of IRS insecticide diversification. Furthermore, even combining IRS micro-mosaics with untreated bednets gives estimates of *Λ_Ω,0_*, again relative to undiversified IRS in the absence any bednets, ranging from only 1.32 for *N* = 2 to 1.42 for *N* = 3 or 4. However, supplementing even blanket coverage of IRS with a single active (*N* = 1) with NG-LLINs yields *Λ_Ω,0_* estimates ranging from 1.77 for nets with excito-repellent properties to 1.97 for those that lack them. Combining NG-LLINs with IRS micro-mosaics, is expected to achieve slightly better *Λ_Ω,0_* values, ranging from 2.23 for excito-repellent nets with only two IRS actives (*N* = 2) to 2.44 for nets with purely lethal active ingredients and three, four or presumably more IRS insecticides (*N* ≥ 3).

This expected doubling or better of the interval to resistance trait emergence against IRS actives, achieved primarily with NG-LLIN use and secondarily with micro-mosaic formats for IRS deployment, may be useful from an IRM perspective but that depends on baseline frequency at which incipient resistance traits occur naturally. For example, a 10-year lifetime for pre-emptive insecticide usage under baseline conditions, as a stand-alone IRS intervention with a single active (*λ_0_*), may be extended to 20 years or more by combining with NG-LLINs and, to a lesser extent, deployment as micro-mosaics alongside complementary IRS actives (*λ_Ω_*). However, insecticides for which incipient resistance traits naturally occur more frequently will have correspondingly shorter extended lifetimes.

Furthermore, these simulations also suggest quite fundamental and substantive constraints upon the upper limits of realistic expectations imposed by behavioural processes that lie beyond the biological reach of indoor vector control interventions (Figures 5, 6 and 7). Although such reductions of indoor survival and feeding success by NG-LLINs are expected to approximately half the overall probability of an *F_0_* mosquito carrying a novel resistance trait from reproducing (*P_ϕ,≤1_*), none of the simulated combinations of NG-LLINs with IRS micro-mosaics are expected to reduce this sparking rate by much more than three fifths, regardless of the diversity of insecticides deployed through the latter measure (Figure 7). Correspondingly, while NG-LLINs alone may be reasonably expected to approximately double timelines to the emergence of resistance into *F_≥1_* generations that subsequently experience selection pressure, supplementing such NG-LLINs with IRS micro-mosaics is expected to delay such events by only 2.5-fold, with negligible incremental benefit achieved IRS active ingredient diversification beyond the minimum level of two insecticides (Figure 7). This is because their impact upon emergence timelines is limited by even the small fractions of zoophagy and exophagy that may occur among predominantly anthropophagic, endophagic mosquitoes like *An. funestus* (Figure 5 and 6).

Specifically, the biggest constraint on this pre-emptive resistance management function of either type of LLINs, regardless of IRS active diversification level, is simply that they don’t protect people or kill mosquitoes outdoors. Under the assumed conditions of the *An. funestus* scenarios considered in this modelling exploration, a consistent fraction of mosquitoes (19%) are expected to feed outdoors during their first feeding attempt without any exposure to LLINs or IRS, irrespective of the diversity of active ingredients used in either vector control measure (Figure 5). This considerable minority of mosquitoes experience no indoor insecticide exposure, so equivalent fractions of incipient resistance alleles occurring among *F_0_* individuals will survive the gauntlet of indoor control with two or more complementary insecticides, regardless of whether they are encountered on an NG-LLIN and/or on an IRS-treated surface. So, while the rates at which the sparks of novel resistance traits survive and progress beyond *F_0_* may be suppressed, correspondingly extending timelines to emergence of *F_≥1_* generation wildfires that will be fanned by the flames of selection for those traits, there are considerable fundamental constraints upon how much may be reasonably expected of such approaches to delaying the inevitable. As for the distinct issue of residual malaria transmission, which occurred ubiquitously even before widespread physiological resistance [31, 32], the limitations of insecticide resistance strategies based solely on indoor vector control measures are primarily defined by the ability of mosquitoes to evade them by feeding outdoors without ever entering a house.

A secondary contributing pathway to persisting resistance emergence beyond *F_0_*, observed across all intervention scenarios involves smaller fractions of mosquitoes that are exposed to a single IRS insecticide indoors but fail to feed successfully and subsequently obtain a blood meal outdoors during a second attempt, again usually outdoors (Figure 5). In this case, mosquitoes avoid sequential exposure to a second IRS insecticide, thereby retaining the capacity to pass on resistance traits into subsequent generations. Although the expected fractions of novel resistance traits expected to become established beyond *F_0_* vary somewhat, depending on NG-LLIN product profile and IRS insecticide diversification, they make a consistently lower overall contribution to delaying their emergence into *F_≥1_* generations than those that simply feed outdoors at the first attempt, sometime on animals.

Given the key role of even occasional zoophagy and exophagy in constraining the impact of these IRM strategies on resistance emergence timelines in such an anthropophagic and endophagic vector as *An. funestus*, it comes as no surprise that equivalent simulations for *An. arabiensis* indicate expected impacts on this species are negligible (Figures 7 and 8). For this behaviourally plastic vector that opportunistically expresses zoophagy and exophagy as readily as it does anthropophagy and endophagy, the effects of all these various intervention combinations on resistance dynamics are entirely negated by the high frequency of these former behaviours (Figure 7). No combination of NG-LLINs and/or IRS micro-mosaics is expected to meaningfully alter either the probability of resistance emergence (*P_ϕ,≤1_*) or the strength of selection pressure acting on resistance traits once they become established (*s*). In the conceptual context of Figure 3 and Box 1, this reflects the fact that rates of spark generation (*P_ϕ,≤1_*) are consistently even higher for *An. arabiensis* than for *An. funestus*, while wildfire intensity (*s*) are consistently intrinsically much lower, not because these interventions lack efficacy but rather because mosquitoes largely avoid exposure to them altogether. For the same underlying reasons, these respective primary and secondary outcomes are expected to be essentially unchanged by any combination of NG-LLINs and/or IRS micro-mosaics (Figure 7).

Indeed, the dominant behavioural feature underpinning this pattern is the high prevalence of outdoor feeding, often on animal hosts, particularly during the first feeding attempt (Figure 8). This results in over 85% of *An. arabiensis* mosquitoes successfully obtain blood outdoors in their first feeding attempt without entering houses, thereby completely avoiding indoor interventions. Consequently, almost all mosquitoes carrying incipient resistance traits experience no insecticide exposure whatsoever, eliminating the opportunity for IRM interventions to suppress the emergence of those traits. On the other hand, these same evasive behaviours are also expected to consistently moderate the evolutionary pressures that favour selection of such resistance traits after they do emerge.

**Figure 8:**
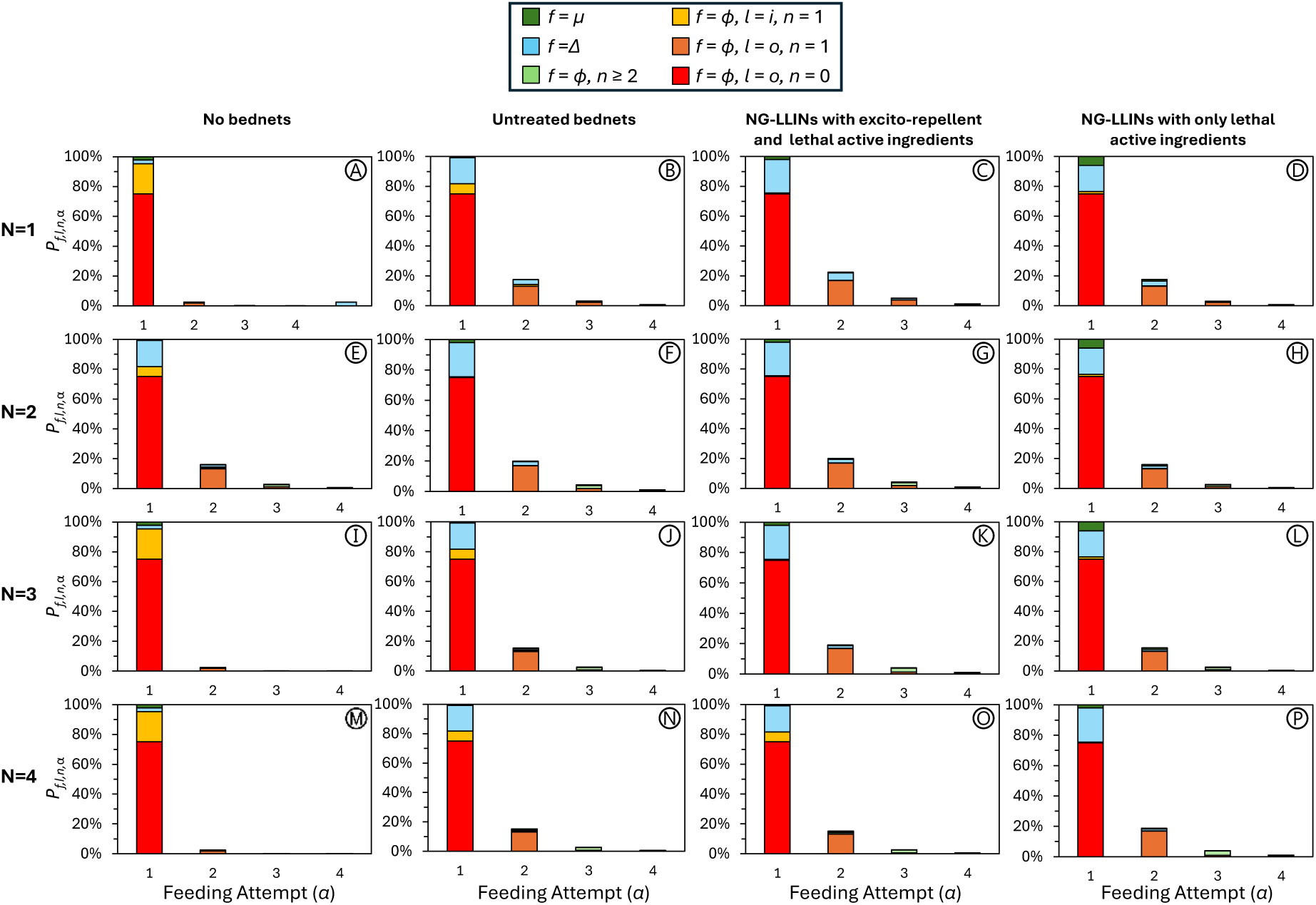
The expected proportions of host-seeking *An. arabiensis* mosquitoes starting a gonotrophic cycle (*P_f,l,n,α_*) that experience a deterred, died or successfully fed fate (*f = Δ*, *μ* or *ϕ*, respectively) at either an indoor or outdoor location (*l* = *i* or *o*, respectively) after exposure to none, one or several of the insecticides used for indoor residual spraying (IRS; *n* = 0, 1 or ≥2) in the first, second, third or fourth feeding attempt (*α* = 1, 2, 3 or 4) for all possible intervention scenarios combining either no bednets, untreated bednets, next generation long-lasting nets (NG-LLINs) exhibiting both lethal and excito-repellent properties or NG-LLINs exhibiting only lethal modes of action with IRS using either blanket coverage with only one insecticide (*N* = 1) or household-scale micro-mosaic formats with two to four insecticides (*N* = 2, 3 or 4).

## Discussion

Taken together, these findings indicate that next-generation long-lasting insecticidal nets (NG-LLINs), particularly those with predominantly lethal modes of action, may usefully delay the emergence of resistance to IRS insecticides in highly anthropophagic and endophagic vectors like *An. funestus*. When combined with IRS micro-mosaics, this delay may be extended somewhat further but it is the NG-LLINs that contribute most to such lengthening of pre-emptive use lifetimes for novel active ingredients. Within the conceptual framework outlined in Figure 3, reductions in resistance emergence operate primarily by lowering the probability that resistant *F₀* mosquitoes survive and reproduce (*P_ϕ,≤1_*), thereby prolonging the delay phase preceding resistance establishment. As such, the delay in resistance emergence observed here reflects the prevention of spontaneously occurring novel resistance traits in individual *F_0_* insects from establishing themselves in *F_1_* and subsequent generations during the early emergence phase, rather than any worthwhile reduction in their survival and reproduction advantage after they become established and are subjected to sustained directional selection over indefinite periods. In terms of the analogy with wildfires, this may be conceptualized as dampening of the frequency that sparks sporadically occur and ignite small flames, rather than appreciable suppression of the rate with those fires subsequently spread across the forest.

A key implication of these findings is that the resistance management value of IRS micro-mosaics depends on widespread and consistent use of NG-LLINs treated with active ingredients that are novel and pre-emptively introduced together, rather than in a reactive one-at-a-time manner that allows vector populations to steadily evolve resistance against each of them sequentially [19, 43]. Without such NG-LLINs, most *An. funestus* mosquitoes are expected to feed successfully indoors on their first attempt after encountering only a single IRS insecticide, regardless of the diversity of insecticides used for IRS. As a result, any such successfully fed *F_0_* individual that carries a new resistance trait against the particular sole IRS active it was exposed to is expected to usually reproduce successfully, before any further exposure. It is therefore unreasonable to expect fine scale diversification of IRS actives through micro-mosaic deployment formats to have any meaningful impact upon the emergence rate of such novel traits unless it is used to supplement LLINs, NG-LLINs in particular, as the primary malaria vector control intervention. Correspondingly, gaps in LLIN availability or use may be expected to undermine the effectiveness of IRS micro-mosaics as supplementary IRM interventions. While this highlights the need for strategies that remain robust under real-world conditions, it is encouraging that high LLIN coverage has been sustained across most sub-Saharan African settings [39, 44], so it may be feasible for control programmes to consider combining these interventions for IRM purposes.

For anthropophagic and endophagic vectors like *An. funestus* [45, 46], NG-LLINs primarily delay resistance through high mosquito mortality arising from exposure to their two complementary active ingredients, the vast majority of which occurs during first and second feeding attempts (Figure 5). Encouragingly, NG-LLINs with predominantly lethal modes of action are predicted to be most effective for delaying the emergence of resistance against IRS actives for the same reasons that they achieve greater overall impact on malaria transmission despite conferring less personal protection than nets with stronger excito-repellent properties: They more frequently kill mosquitoes outright, rather than merely force them to look elsewhere for blood, so no trade-offs need to be made when selecting net products to both maximize immediate epidemiological impact and long-term effectiveness as IRM tools [47-49].

Despite the transformative effects of NG-LLINs on the predicted life histories of this hypothetical *An. funestus* population, specifically because the predominantly anthropophagic and endophagic behaviours assumed render it highly vulnerable to such an indoor-targeted vector control measure, a noteworthy fraction of mosquitoes are nevertheless expected to feed and then go on to successfully reproduce. Even when fully diversified (*N* = 4) IRS micro-mosaics are deployed, between a fifth (No bednets) and a third (Excito-repellent NG-LLINs) are expected to successfully feed outdoors after exposure to only one IRS active ingredient or none at all. These sizable minority fractions of fates that ultimately result in successful feeding therefore maintain a minimum baseline risk of resistance emergence that indoor vector control simply cannot eliminate for fundamental behavioural reasons [32], even for such an anthropophagic, endophagic vector as the stereotypical *An. funestus* population simulated here.

By corollary, this limitation is of course far greater for more exophagic and zoophagic species like *An. arabiensis* (Figure 6,7 and 8), for the same reasons that they are often important contributors to persistent residual malaria transmission [27, 31, 32, 50]: They largely avoid indoor exposure to NG-LLINs and IRS altogether by feeding outdoors, sometimes on people and sometime on other mammals (Figure 4). This highlights the key role of such evasive behaviours in robustly sustaining both stable residual malaria transmission [27, 31, 32, 50] and risks of novel resistance trait emergence. Note, however, that such zoophagic and exophagic behaviours may attenuate the potential consequences of resistance emergence over the long term because they may be described as behavioural refugia against insecticide exposure, thus limiting selection pressure favouring any resistance traits that do emerge [51]. Indeed, this principle is fully reflected in the far lower selection coefficients calculated herein for *An. arabiensis* (Figure 7) and empirical evidence from all across Africa [52, 53]. Nevertheless, IRS micro-mosaics are expected to provide negligible IRM benefit for *An. arabiensis*, regardless of how well diversified IRS insecticides are at household scale (Figure 7), simply because they primarily feed outdoors, often upon animals (Figures 6 and 8).

Furthermore, even for vectors like *An. funestus* that predominantly feed indoors upon humans, the availability of occasional outdoor feeding opportunities also limits the value of diversifying IRS actives beyond two insecticides in any given place at any given time. The limited impact upon insecticide resistance emergence trajectories predicted for *An. funestus* reflects the expectation that, across all simulated intervention scenarios, the vast majority of such mosquitoes that survive after failing to feed in their first attempt subsequently die or divert to outdoor feeding on their second attempt. Even for such a specialized, human-dependent mosquito such as *An. funestus*, the limited behavioural plasticity they exhibit may nevertheless facilitate a diversity of very different life histories that all almost always conclude with either successful blood acquisition or death within one or two feeding attempts (Figure 5), leaving very few individuals surviving long enough to experience repeated encounters with multiple insecticides across subsequent feeding attempts. Consequently, the intended resistance management benefit of IRS micro-mosaics through sequential exposure to diverse insecticides may only be partially realised in practice, because mosquitoes are only rarely exposed to more than two IRS insecticides before they either feed or die.

The diminishing returns obtained from diversifying IRS micro-mosaics beyond two insecticides in a given time and place must therefore be carefully weighed against the increasing operational complexity and cost associated with implementing multi-insecticide IRS strategies, including procurement, logistics, training, and safe disposal [54]. A more efficient approach may therefore be to deploy a broader portfolio of complementary active ingredients more dynamically and heterogeneously. For example, programmes could implement household-scale micro-mosaics with two distinct IRS insecticides in any given year, while rotating between two or more different insecticide pairs on an annual basis [22, 55]. And individual provinces, districts, counties or other administrative unit within a country could be assigned to different pairs of active ingredients at any given time and rotated through the alternative pairs in different orders, so that all four or more formulations may be procured steadily in approximately similar quantities each year. This in turn would make planning and resourcing far easier by stabilizing budget requirements at national scale, while simultaneously maximizing insecticide deployment heterogeneity across space and time at local scales. Such an approach would therefore balance short term practicality with long term effectiveness for IRM purposes without introducing significant financial or operational complexity.

The appropriateness of NG-LLINs and supplementary IRS micro-mosaics may be highly context dependent, so their deployment must be guided by local vector ecology. In settings characterised by low transmission intensity and predominantly zoophagic, outdoor-feeding mosquito populations, such as parts of the Americas [31, 32, 56, 57], indoor-focused interventions may achieve limited impact [27, 32, 58] and, consistent with the results of one of the few controlled IRM trials conducted in such circumstances [59, 60], resistance evolution trajectories may be largely unaffected by insecticide deployment regime (Figure 7). In contrast, in high transmission regions such as sub-Saharan Africa and parts of Oceania, where human-specialised, predominantly indoor-feeding vectors have historically mediated most transmission [31, 32, 56, 57], NG-LLINs and IRS micro-mosaics may provide greater IRM value. These behavioural differences also have important geographical implications, reinforcing the need to move away from “one-size-fits-all” resistance management strategies and toward approaches stratified by vector behaviour and ecology, often ideally targeting both indoor and outdoor transmission pathways [61, 62].

Note, however, that maximizing impact upon the most efficient anthropophagic and endophagic vectors should always be prioritized. In turn, that may often necessitate trade-offs because the optimal characteristics of vector control measures may vary according to the behavioural characteristics of individual mosquito species within a malaria vector guild. For example, while maximizing excito-repellency and personal protection may optimize impact upon transmission by zoophagic vectors, it may undermine all-important community-level impacts upon transmission by more efficient anthropophagic vectors through population suppression [47-49] and even elimination [30]

Ultimately, improving malaria vector control will not only require optimization of indoor insecticide delivery but also new interventions that address persisting gaps in outdoor protection [50]. Outdoor interventions that actually kill mosquitoes, rather than merely deter them, could not only enhance immediate transmission control [48, 49] but also, as illustrated by these simulations, long-term IRM effectiveness. However, this approach will depend on the development of novel non-pyrethroid insecticides suitable for outdoor use, including vapour-phase formulations for protecting humans while they are awake and active [47, 63-65] and endectocides for targeting mosquitoes when they feed upon livestock [66]. While developing such new interventions will obviously be challenging and expensive, closing such biological coverage gaps [32, 67] could enable unprecedented progress towards malaria elimination, especially in settings dominated by highly anthropophagic vectors that could well be eliminated across a much broader geographic range than they have in the past [30, 68].

Further refinement of the micro-mosaic concept could involve spraying different insecticides to different specific indoor surfaces. However, practical and financial constraints, particularly the high costs associated with spraying large indoor surfaces, remain significant barriers to IRS scale up generally [69], so perhaps the lessons learned from these simulations may be more practically useful in relation to alternative formats for achieving the same IRM goals. For example, perhaps the most obvious lesson learned here is simply that NG-LLINs treated with pre-emptively combined combinations of novel insecticides actually do most of the “heavy lifting” for IRM purposes because they not only deliver those complementary actives in a compact enough format to enable use of the full-dose mixtures that theoretical analysis suggests are optimal [22, 43, 70-72], they also bring the micro-mosaic paradigm down to sub-household scale when combined with IRS inside individual dwellings. With respect to building on the successes of NG-LLINs as a deployment format for full-dose mixtures, an alternative that extends this paradigm to protected entire houses is durable insecticide-treated mosquito-proofed netting that can be readily installed on windows, doors, and eave openings [73-75]. This approach allows re-purposing of IRS formulations, some of which could never be safely deployed in LLIN formats, by enabling application by brush rather than spray to surfaces that protected occupants make far less direct contact with, while also dramatically reducing the total surface area requiring treatment compared to IRS [73, 74]. Such insecticidal mosquito proofing of houses could also enable micro-mosaic deployment of different insecticides, or even different full dose mixtures at sub-household scale, simply by screening different ventilation openings in each house with different pre-treated netting materials and/or retreating them with different complementary formulations. Such practical, affordable new formats for achieving sub-household micro-mosaic formats could force mosquitoes into extended sequences of exposures to two or more insecticides, even while foraging around a single protected house, potentially offering a practical and affordable route to achieving the IRM benefits expected of insecticide combinations and full-dose mixtures [22, 24, 55].

However, several limitations should be considered when interpreting these findings. First of all, this analysis assumes consistent intervention coverage over time and does not incorporate temporal variations in long-lasting insecticidal net (LLIN) usage or insecticide decay. In addition, the model assumes full population coverage with both NG-LLINs and IRS in scenarios where these interventions are deployed. In reality, this goal is never more that approximately approached, so the insecticide resistance management benefits predicted herein may not be fully realized in practice.

Second, the model framework assumes a fully pre-emptive strategy in which all insecticides used for NG-LLINs and IRS are novel and introduced simultaneously, prior to the emergence of relevant resistance traits. This scenario may be unrealistic under current conditions, due to new chemical classes for IRS being prohibitively expensive for routine programmatic deployment in ideal formats.

Third, while the analysis focuses on resistance dynamics associated with IRS active ingredients, it does not explicitly model resistance evolution to NG-LLIN actives. This relatively simple approach may therefore overlook potential interactions between resistance mechanisms across intervention types.

Fourth, the model adopts a deliberately simple deterministic structure, generating single value estimates of feeding cycle outcomes rather than incorporating stochastic variability. It also does not explicitly project resistance allele frequency trajectories over time. This limits the ability to capture random effects, heterogeneity, and complex evolutionary dynamics that may arise in real populations.

Taken together, these limitations highlight that model outputs should be interpreted with caution when extrapolating to field settings, where ecological, behavioural, and operational complexities may substantially alter transmission dynamics and resistance outcomes.

## Conclusion

Despite these limitations, this modelling analysis nonetheless facilitates several practically useful insights. Overall, NG-LLINs and, to a lesser extent, IRS micro-mosaics may usefully delay the emergence of novel insecticide resistance traits, but these IRM benefits are intrinsically dependent upon strong vector behaviour preferences for feeding indoors upon humans. The constraints imposed upon reasonable expectations of IRM benefits by vectors that feeding outdoors are modest but nevertheless important even for mosquitoes like *An. funestus* that only do so occasionally. Correspondingly, no meaningful IRM effects may be reasonably expected for predominantly outdoor-feeding vectors such as *An. arabiensis*, although these species represent only secondary targets for control wherever transmission is dominated by more efficient, human-specialized vectors that usually feed mostly indoors.

Regarding the predominantly anthropophagic and endophagic malaria vectors such interventions are most appropriate for, they may suppress the initial survival and reproduction of individuals carrying new resistance traits against IRS insecticides, thereby delaying the emergence of those traits by 2 to 2.5-fold. However, they cannot be reasonably expected to delay such inevitable events much more than that, or to sufficiently reduce selection pressures acting on their progeny to stifle their subsequent spread throughout the population. Achieving durable vector control and sustainable resistance management will therefore require integrating these tools with complementary strategies that address outdoor transmission and reduce the ecological opportunities for both resistance emergence and spread.

## Data Availability

The data used in this analysis has been uploaded as a Supplementary File 2.

## Abbreviations

GPIRM: global plan for insecticide resistance management
IRS: indoor residual spraying
LLIN: long lasting insecticidal treated nets
PBO: piperonyl butoxide
NG-LLINs: next generation long lasting insecticidal nets.

