## Supplementary File 1 for "Potential impacts of supplementing next generation long-lasting insecticidal nets with household-scale micro-mosaic deployment of indoor residual spraying with insecticides upon rates of incipient resistance trait emergence and selection"

### Potential impacts of supplementing next generation long-lasting insecticidal nets with household-scale micro-mosaic deployment of indoor residual spraying with insecticides upon rates of incipient resistance trait emergence and selection

**Dingani Chinula[1,2], Nicholaus Mziray[3], Neil Philip Hobbs[4], Busiku Hamainza[1], Thomas Reed[2,5], Samson Kiware[3] and Gerry F. Killeen[2,5]**

1. National Malaria Elimination Centre, Chainama Hills Hospital Grounds, PO Box 32509, Lusaka, Republic of Zambia
2. School of Biological, Earth and Environmental Sciences, University College Cork, Cork, Republic of Ireland
3. Department of Environmental Health and Ecological Sciences, Ifakara Health Institute, Ifakara, Morogoro, United Republic of Tanzania
4. Department of Vector Biology, Liverpool School of Tropical Medicine, Liverpool, United Kingdom.
5. Sustainability Institute, University College Cork, Cork, Republic of Ireland

#### Introduction

Here we present a novel deterministic mathematical model for assessing mosquito life histories as a function of their host-seeking behaviours, behavioural responsiveness to excito-repellent active ingredients and insecticide susceptibility traits. This model is then applied to exploring implications for IRM in the context of a malaria vector control scenario in which universal coverage of NG-LLINs is supplemented with IRS applied as a micro-mosaic of complementary active ingredients at household scale (*Introduction* and outline *Methods* sections in the main manuscript).

#### Detailed Methods

##### Mathematical notation and definitions

For clarity and ease of reference, all the model parameters are defined in Table S1 and all the equations required are presented in Table S2. For ease of comparison, the symbols and definitions used herein are as consistent as possible with those of previous modelling studies by some of the same authors [1-5] and illustrated intuitively in Figure S1. In the interest of clarity and parsimony, a number of simplifying assumptions were made, which are explained and justified as follows.

##### Simplifying assumptions regarding mosquito biology, life history and intervention characteristics

In the conceptual framework of this model (Figure 4 in the main manuscript), it is assumed that mosquitoes feed upon animals or humans, do the latter either indoors (*i*) or outdoors (*o*), and may be either deterred ($\Delta$), killed ($\mu$) or feed successfully ($\phi$). The latter process of successful feeding and subsequent survival until oviposition is completed, is even assumed to sometimes happen inside houses where humans are well but imperfectly protected by NG-LLINs. However, although never entirely true in reality, it is assumed for the sake of simplicity that entering a house is inevitably associated with exposure to any IRS insecticide that has been applied therein. It is also assumed that mosquitoes may instead feed outdoors upon unprotected animals or humans, which may often be preceded by exposure to one or more IRS insecticides during unsuccessful previous feeding attempts inside houses, during which no unprotected host outside of an NG-LLIN could be successfully accessed.

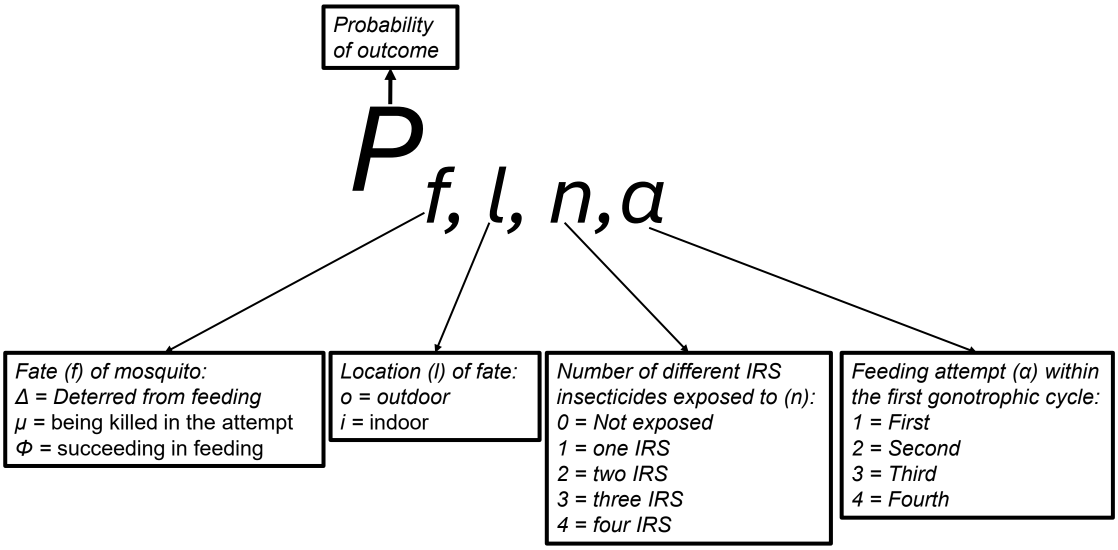


**Figure S1. A schematic illustration of the notation used herein to convey the meaning of all the different conditional probabilities calculated in the following deterministic model of mosquito life histories.**

An important simplifying assumption made is that the first gonotrophic cycle is the most important one for resistance propagation because it is inevitably the most commonly completed one, even in quite long-lived populations, and because the cumulative probabilities of a mosquito completing subsequent gonotrophic cycles approximate to a compound function of that first one.

It is assumed, however, that the first gonotrophic feeding cycle can be broken down into sequential discrete feeding attempts ($\alpha$) undertaken until the mosquito either succeeds in feeding ($\phi$) or dies in the attempt ($\mu$). It is also assumed that a mosquito may make up to four sequential attempts to feed ($\alpha$ = 1, 2, 3 or 4), with the probabilities of completing four or more feeding attempts ($\alpha \geq 4$) without success assumed to be negligible *a priori* ($P_{\Delta,i,4} \approx 0$).

Other simplifying assumptions regarding mosquito life history include lack of gonotrophic discordance, notably the pre-gravid feeding that occurs so frequently among freshly emerged females in African *Anopheles* populations [6-11]. This assumption, although clearly inaccurate in many cases, is justified on the basis that females taking partial pre-gravid bloodmeals cannot oviposit without needing a subsequent full bloodmeal [6-11]. It is therefore assumed those that do so indoors will usually remain within a single house over the course of both these feeding and resting events.

#### Table S1: Key model parameter symbols and description

| Notation | Definition and explanation |
|---|---|
| *Input parameters* | |
| $Q_h$ | Proportion of mosquitoes that would attempt to feed upon humans (*h*) rather than animals, estimated as the proportion that obtain a blood meal from humans [12, 13] under a baseline condition lacking any NG-LLINs or IRS. Note that this term has been annotated a little more explicitly as $Q_{h,0}$ in some similar models previously used to explore the roles of insecticidal person protection measures in malaria transmission control [3, 14], wherein $Q_{h,\Psi}$ described $Q_h$ under a given intervention scenario ($\Psi$). Here, however, the annotation of this parameter is simplified to accommodate all the other subscripts required to denote the various new details of gonotrophic cycle history and outcomes that are considered. |
| $\pi_i$ | Proportion of mosquitoes attempting to feed upon humans that do so by entering a house to forage indoors (*i*), estimated as the proportion of attacks upon humans that occur indoors in the absence of LLINs, IRS, mosquito-proofed housing, repellents or any other personal protection measure, based on representative measurements of behavioural interactions between mosquitoes and humans under such baseline conditions [15, 16]. |
| $\Delta$ | Proportion of mosquitoes deterred from feeding on humans indoors before they can experience fatal exposure to active ingredients of any NG-LLINs present. Note that this overall proportion encompasses all possible causes for the mosquito to abandon that feeding attempt, including human host defensiveness, the protective physical barriers represented by bednets or by excito-repellent insecticides used to treat those nets and turn them into NG-LLINs, that occurs soon enough to prevent fatal exposure to the latter [3, 17, 18] consistent with standard WHO definitions [19]. |
| $M$ | Proportion of mosquitoes that die as a result of attempting to feed on a human host, either through natural mortality processes or through contact with the active ingredients of NG-LLINs, regardless of whether they die before or after feeding [3, 17, 18] as per standard WHO definitions [19]. |
| $N$ | Total number of distinct insecticides formulations used for IRS as either singular blanket coverage ($N$ = 1) or a household scale micro-mosaic format ($N$ > 1). |
| *Foraging history stages, intermediate fates and final fates at different locations* | |
| $A$ | Feeding attempt in the first gonotrophic cycle ($\alpha$ = 1, 2, 3 or 4; See assumptions for limiting to ≤4) |
| $N$ | Total number of different IRS insecticides that a mosquito has been exposed to inside one or more houses over the course of a given gonotrophic cycle ($n$ = 0, 1, 2, 3 … $N$), categorized as $n$ = 0,1 or ≥ 2. |
| $F$ | Intermediate or final fate, classified in terms of either of the three following possible outcomes deterrence(Δ), mortality ($\mu$)or successful feeding ($\phi$). |
| $\Delta$ | Intermediate fate ($f$) of deterrence, meaning that the mosquito abandoned a given feeding attempt ($\alpha$) before it could successfully feed or be killed by natural mortality factors or the active ingredients of NG-LLINs. |
| $\Phi$ | Final fate ($f$) of having eventually succeeded in feeding and reproducing by laying eggs. |
| $M$ | Final fate ($f$) of having died in the attempt to feed. |
| $L$ | Location at which a given intermediate or final fate occurred, classified in terms of the following two broad types of environments. |
| $I$ | Indoor location ($l$) at which a given intermediate or final fate occurs |
| $O$ | Outdoor location ($l$) at which a given intermediate or final fate occurs |
| *Intermediate and final output parameters* | |
| $P_{f,l,n,\alpha}$ | Probability of a mosquito experiencing a given fate in the first gonotrophic cycle ($f = \Delta$, $\mu$ or $\phi$) in a given location ($l$ = $i$ or $o$) after exposure to $n$ IRS micro-mosaic insecticides over $\alpha$ feeding attempts ($\alpha$ = 1, 2, 3 or 4). Refer upper portion of Figure S1 for clear representation of the meaning of this quite complex set of four parameters in the subscripts, each representing two to four distinct possible conditions. |
| $P_{\phi,o,n,\alpha}$ | Probability of mosquitoes successfully feeding in the first gonotrophic cycle ($f = \phi$) outdoors ($l = o$) after exposure to $n$ IRS micro-mosaic insecticides over $\alpha$ feeding attempts ($\alpha$ = 1, 2, 3 or 4). |
| $P_{\phi,i,n,\alpha}$ | Probability of mosquitoes successfully feeding in the first gonotrophic cycle ($f = \phi$) indoors ($l$ = i) after exposure to $n$ IRS micro-mosaic insecticides over $\alpha$ feeding attempts ($\alpha$ = 1, 2, 3 or 4). |
| $P_{\Delta,i,n,\alpha}$ | Probability of mosquitoes being deterred in the first gonotrophic cycle ($f = \Delta$) indoors ($l = i$) after exposure to $n$ IRS micro-mosaic insecticides over $\alpha$ feeding attempts ($\alpha$ = 1, 2, 3 or 4). |
| $P_{\mu,i,\alpha}$ | Probability of mosquitoes dying in the first gonotrophic cycle ($f = \mu$) indoors ($l = i$) after exposure to NG-LLINs and $\alpha$ feeding attempts ($\alpha$ = 1, 2, 3 or 4), regardless of the number of IRS insecticides encountered. |
| $P_{\phi,\leq 1}$ | **Primary outcome:** Probability of mosquitoes that successfully feed and survive encounters with NG-LLINs and other indoor hazards during their first gonotrophic cycle after having been exposed to one, and only one, IRS micro-mosaic insecticide ($n$ = 1) or none ($n$ = 0). |
| $P_{\phi,0}$ or $P_{\phi,o,1,0}$ | Probability of mosquitoes that successfully feeding ($f = \phi$) without indoor exposure to any IRS micro-mosaic insecticide ($n$ = 0), in the first feeding attempt ($\alpha$ = 1) , which can only occur outdoors ($l = o$) as per figure 4 in the main manuscript. |
| $\lambda_\Omega$ | Expected mean lifetime of pre-emptive use for a new IRS insecticide in a given intervention scenario ($\Omega$, where $\Omega = A, B, C$ etc), meaning the lean length of the period over which it can be used in a vector control |

| | |
|---|---|
| | intervention before an incipient resistance trait arises de novo in a single $F_0$ mosquito and is allowed to survive and reproduce, thus establishing itself as a lineage in the population that may be selected for by insecticide pressure from $F_1$ onwards. |
| *A* or *B* | Identifiers for two distinct vector control intervention scenarios to be compared and contrasted in terms of expected mean lifetime of pre-emptive use for a new IRS insecticide ($\lambda_A$ and $\lambda_B$ , respectively). |
| $\Lambda_{B,A}$ | Relative duration of the expected mean lifetime for pre-emptive use for an insecticide in one pre-emptive intervention scenario versus another ($\lambda_\Omega$ for scenario *B* relative to scenario *A*). |
| *S* | **Secondary outcome**: Evolutionary *selection coefficient* [20, 21] favouring mosquitoes with resistance traits against one of the IRS insecticides, calculated as the fraction of mosquitoes surviving all feeding attempts required to complete their first blood meal that have been exposed to one and only one IRS micro-mosaic insecticide. |

A number of assumptions are also made regarding the insecticidal active ingredients deployed in NG-LLIN (designated *x* and *y*) and IRS formulations (designated *a*, *b*, *c*, etc.), the most important being that they are all complementary to each other with no overlapping mechanisms of action or resistance traits. Also, while the physical barrier and excito-repellent actives of the NG-LLINs may deter mosquitoes from attacking their human occupants ($\Delta$), the IRS insecticides are assumed to lack any repellent or irritant properties, only inducing mortality through direct physical contact.

It is also assumed that while the active ingredients of NG-LLINs are fast-acting, those used for the IRS micro-mosaics act much more slowly. Herein, NG-LLINs are assumed to kill mosquitoes either immediately before they feed or within hours of doing so, leaving them no opportunity to subsequently gestate and oviposit and preventing successful completion of the first gonotrophic cycle. IRS with even very slow-acting active ingredients can have impressive impacts upon malaria vector populations and the transmission they mediate because the sporogonic incubation period of the parasite typically spans several gonotrophic cycles, involving multiple visits to different houses by anthropophagic vector species [3, 22]. It is therefore assumed that while IRS exposure kills mosquitoes before they complete gestation and then oviposit, this takes one or two full nights, so that the number of different IRS actives cumulatively encountered over repeated visits to houses over several feeding attempts during that period can be simply calculated without having to explicitly account for IRS-related mortality.

**Possible outcomes of the first feeding attempt by a freshly emerged mosquito**

Upon emerging from a breeding habitat, an adult female mosquito seeks out a first blood meal to support egg production and complete its first gonotrophic cycle (Figure 4 in the main manuscript). Assuming that contact with IRS or NG-LLIN insecticides only occurs inside houses, that depends upon the probabilities of both attacking a human ($Q_h$) and entering a house to do so ($\pi_i$). The parameter $Q_h$, often referred to by entomologists as the human blood index [12, 13], is defined as the proportion of bloodmeals that would be obtained from humans in the absence of any intervention, just as in previous models [3, 23] (Table S1). For this application $\pi_i$ is defined slightly differently as the proportion of mosquitoes attacking humans that do so indoors, although it is nevertheless measured in the same way [15, 16] (Table S1). In practical terms, $\pi_i$ is measured as the baseline proportion of human exposure that happens indoors in the absence of IRS, LLINs, mosquito-proofed housing, repellents or any other protective measure against mosquito bites [15, 16]. This summary measure is estimated by measuring indoor and outdoor biting rates on unprotected humans over each hour of the night and weighting these otherwise misrepresentative exposure rates according to the proportions of people who report being indoors or outdoors at those times [15, 16] . This approach captures the dynamic interaction between mosquito activity and human behaviour throughout the night, providing a realistic measure of not only indoor exposure risk under baseline conditions but also the potential effectiveness of indoor-targeted interventions against malaria transmission like LLINs and IRS [15, 16, 24, 25]. While this rephrasing does not alter the meaning of $\pi_i$ or how it is measured in practice [15, 16] it does more readily allow the underlying principles of this model to be conceptualized.

However, not all mosquitoes enter houses (Figure 4 in the main manuscript) and many may avoid exposure to NG-LLINs and IRS by either feeding outdoors upon non-human animal hosts or upon unprotected humans (Table S2, Eq. 1.1). Such mosquitoes are assumed to feed successfully ($f = \phi$), gestate and subsequently lay eggs as no IRS or LLIN exposure effects apply. Such zoophagic and exophagic tendencies represent a behavioural refuge against insecticide selection pressures that simultaneously undermines intervention impact while also attenuating selection pressure for resistance. Hence, we have included green shading around Eq. 1.1 in Table S2, reflecting the lack of any prior foraging visits inside houses ($n$ = 0 out of $N$) for mosquitoes that find and obtain blood from unprotected humans or animals outdoors ($l = o$) on their first feeding attempt ($\alpha$ = 1).

**Table S2. Equations for calculating the probabilities (*P*) of each of seven possible combinations of the distinct possible fates of mosquitoes (*f*) in either indoor or outdoor locations (*l*), resulting in cumulative exposed to given total numbers (*n*) of all the different insecticides deployed as IRS micro-mosaics (*N*), over the first four possible feeding attempts ($\alpha$ = 1 ,2, 3 or 4) of their first gonotrophic cycle ($P_{f,l,n,\alpha}$).** The table cell shaded in red represent the fractions of mosquitoes that are not exposed to the active ingredients of either indoor residual spraying (IRS) or next generation long-lasting insecticidal net (NG-LLIN) over their first gonotrophic cycle because they fed outdoors upon unprotected animals or humans on their first feeding attempt without ever visiting a house. Table cells with blue shading represent the fractions of mosquitoes that have been inside at least one house and exposed to at least one of the IRS insecticides deployed as a micro-mosaic but have yet to feed. Table cells with orange or brown shading represent the fractions of mosquitoes that have been exposed to only one IRS insecticide inside one or more houses and have successfully fed either indoors (dark orange) or outdoors (brown), allowing those carrying incipient resistance traits against that single IRS insecticide to propagate into subsequent generations. Table cells with green shading represent the fractions of mosquitoes that do not contribute to further propagation of any resistance traits they may carry against one IRS insecticide (*a*), because they were killed quickly by either natural hazards or the NG-LLIN active ingredients (*x* and *y*; dark green shading) or more slowly later following exposure to two or more IRS insecticides (light green shading).

| Mosquito Fate (*f*), location (*l*) and cumulative IRS insecticide diversity exposure history (*n*) | First feeding attempt ($\alpha$ = 1) | Second feeding attempt ($\alpha$ = 2) | Third feeding attempt ($\alpha$ = 3) | Fourth feeding attempt ($\alpha$ = 4) |
|---|---|---|---|---|
| Successfully fed ($f=\phi$) upon an unprotected animal or human outdoors (*o*) after exposure to given number (*n*= 0, 1 or ≥2) of all the different IRS insecticides deployed as a micro-mosaic (*N*) after a given number (α) of feeding attempts ($P_{\phi,o,n,\alpha}$) | **Eq. 1.1** $P_{\phi,o,0,1} = 1\text{-}Q_h\pi_i$ | **Eq. 2.1** $P_{\phi,o,1,2} = P_{\Delta,i,1,1}\,(1\text{-}Q_h\pi_i)$ | **Eq. 3.1.1** $P_{\phi,o,1,3} = P_{\Delta,i,1,2}\,(1\text{-}Q_h\pi_i)$<br>**Eq. 3.1.2** $P_{\phi,o,\geq 2,3} = P_{\Delta,i,\geq 2,2}\,(1\text{-}Q_h\pi_i)$ | **Eq. 4.1.1** $P_{\phi,o,1,4} = P_{\Delta,i,1,3}\,(1\text{-}Q_h\pi_i)$<br>**Eq. 4.1.2** $P_{\phi,o,\geq 2,4} = P_{\Delta,i,\geq 2,3}\,(1\text{-}Q_h\pi_i)$ |
| Entered a house but was deterred from attacking any of the occupants inside by the next generation long-lasting insecticidal nets (NG-LLINs) they were using, resulting in the mosquito leaving the house and beginning a subsequent attempt to feed after being exposed to a given number (*n*= 1 or ≥2) of all the different IRS insecticides deployed as a micro-mosaic (*N*) after a given number (α) of feeding attempts ($P_{\Delta,i,n,\alpha}$). | **Eq. 1.2** $P_{\Delta,i,1,1} = Q_h\pi_i\Delta$ | **Eq. 2.2.1** $P_{\Delta,i,1,2} = P_{\Delta,i,1,1}\,Q_h\,\pi_i\,\Delta\,/\,N$<br>**Eq. 2.2.2** $P_{\Delta,i,\geq 2,2} = P_{\Delta,i,1,1}\,Q_h\,\pi_i\,\Delta\,(N\text{-}1)\,/\,N$ | **Eq. 3.2.1** $P_{\Delta,i,1,3} = P_{\Delta,i,1,2}\,Q_h\pi_i\Delta\,/\,N$<br>**Eq. 3.2.2** $P_{\Delta,i,\geq 2,3} = ((P_{\Delta,i,1,2}\,(N\text{-}1)\,/\,N) + P_{\Delta,i,\geq 2,2})\,Q_h\pi_i\Delta$ | **Eq. 4.2.1** $P_{\Delta,i,1,4} = P_{\Delta,i,1,3}\,Q_h\pi_i\Delta\,/\,N$<br>**Eq. 4.2.2** $P_{\Delta,i,\geq 2,4} = ((P_{\Delta,i,1,3}\,(N\text{-}1)\,/\,N) + P_{\Delta,i,\geq 2,3})\,Q_h\,\pi_i\,\Delta$ |
| Entered a house and attacked one of the occupants inside (*l*=*i*) but was killed by the NG-LLINs they were using ($f=\mu$) after a given number ($\alpha$) of feeding attempts ($P_{\mu,i,\alpha}$), regardless of how many (*n*) of all the different IRS micro-mosaic insecticides (*N*) they had been cumulatively exposed to | **Eq. 1.3** $P_{\mu,i,1,1} = Q_h\pi_i\,(1\text{-}\Delta)\,\mu$ | **Eq. 2.3** $P_{\mu,i,1,2} = P_{\Delta,i,1,1}\,Q_h\pi_i\,(1\text{-}\Delta)\,\mu$ | **Eq. 3.3** $P_{\mu,i,1,3} = (P_{\Delta,i,1,2} + P_{\Delta,i,\geq 2,2})\,Q_h\pi_i\,(1\text{-}\Delta)\,\mu$ | **Eq. 4.3** $P_{\mu,i,1,4} = (P_{\Delta,i,1,3} + P_{\Delta,i,\geq 2,3})\,Q_h\pi_i\,(1\text{-}\Delta)\,\mu$ |

| | | | | |
|---|---|---|---|---|
| after a given number (*α*) of feeding attempts ($P_{\mu,i,\alpha}$). | | | | |
| Entered a house and successfully fed (*ϕ*) inside (*l*=*i*), resulting in cumulative exposure to a given number (*n* = 1 or ≥2) of all the different IRS insecticides deployed as a micro-mosaic (*N*). | **Eq. 1.4**<br>$P_{\phi,i,1,1} = Q_h \pi_i (1\text{-}\Delta)\ (1\text{-}\mu)$ | **Eq. 2.4.1**<br>$P_{\phi,i,1,2} = P_{\Delta,i,1,1}\ Q_h \pi_i (1\text{-}\Delta)\ (1\text{-}\mu)\ /\ N$<br>**Eq. 2.4.2**<br>$P_{\phi,i,\geq 2,2} = P_{\Delta,i,1,1}\ Q_h \pi_i (1\text{-}\Delta)\ (1\text{-}\mu)\ (N\text{-}1)\ /\ N$ | **Eq. 3.4.1**<br>$P_{\phi,i,1,3} = P_{\Delta,i,1,2}\ Q_h \pi_i (1\text{-}\Delta)\ (1\text{-}\mu)\ /\ N$<br>**Eq. 3.4.2**<br>$P_{\phi,i,\geq 2,3} = ((P_{\Delta,i,1,2}\ (\text{N-1})\ /\ \text{N}) + P_{\Delta,i,\geq 2,2})\ Q_h \pi_i (1\text{-}\Delta)\ (1\text{-}\mu)$ | **Eq. 4.4.1**<br>$P_{\phi,i,1,4} = P_{\Delta,i,1,3}\ Q_h \pi_i (1\text{-}\Delta)\ (1\text{-}\mu)\ /\ N$<br>**Eq. 4.4.2**<br>$P_{\phi,i,\geq 2,4} = ((P_{\Delta,i,1,3}\ (N\text{-}1)\ /\ N) + P_{\Delta,i,\geq 2,3})\ Q_h \pi_i (1\text{-}\Delta)\ (1\text{-}\mu)$ |
| Expected equalities used to numerically validate each column of feeding attempt specific equations above. | **Eq. 1.5**<br>$P_{\phi,o,0,1}+P_{\Delta,i,1,1}+P_{\mu,i,1}+P_{\phi,i,1,1}=1$ | **Eq 2.5**<br>$P_{\phi,o,1,2}{}^{+}P_{\Delta,i,1,2}+P_{\Delta,i,\geq 2,2}+$<br>$P_{\mu,i,2}+ P_{\phi,i,1,2}+P_{\phi,i,\geq 2,2}$<br>$= P_{\Delta,i,1,1}$ | **Eq 3.5**<br>$P_{\phi,o,1,3}+P_{\phi,o,\geq 2,3}+P_{\Delta,i,1,3}+P_{\Delta,i,\geq 2,3}+P_{\mu,i,1,3}+P_{\phi,i,1,3}+P_{\phi,i,\geq 2,3} =$<br>$P_{\Delta,i,1,2}+ P_{\Delta,i,\geq 2,2}$ | **Eq. 4.5**<br>$P_{\phi,o,1,4}+P_{\phi,o,\geq 2,4}+P_{\Delta,i,1,4}+P_{\Delta,i,\geq 2,4}+P_{\mu,i,1,4}+P_{\phi,i,1,4}+P_{\phi,i,\geq 2,4}$<br>$= P_{\Delta,i,1,3}+P_{\Delta,i,\geq 2,3}$ |
| Primary outcome summarized over all four feeding attempts, namely cumulative proportion of mosquitoes that successfully feed and survive encounters with NG-LLINs indoors after having been exposed to one, and only one, IRS insecticide. | **Eq. 6**<br>$P_{\phi,\leq 1} = P_{\phi,1}+ P_{\phi,0}$<br>$= P_{\phi,o,0,1}+P_{\phi,o,1,2}+ P_{\phi,o,1,3}+ P_{\phi,o,1,4} +P_{\phi,i,1,1}+ P_{\phi,i,1,2} +P_{\phi,i,1,3} +P_{\phi,i,1,4}$ | | | |
| Reciprocal expression of the primary outcome ($P_{\phi,\leq 1}$) in relative terms, to compare two different intervention scenarios (*Ω* = A versus B) in terms of the relative duration of the pre-emptive use lifetime of an insecticide ($\Lambda_{A,B}$), calculated as the quotient of the two relevant pre-emptive use lifetimes in those scenarios ($\lambda_\Omega$). | **Eq 7.1**<br>$\varepsilon_\Omega \propto P_{\phi,\leq 1,\Omega}$<br>**Eq 7.2**<br>$\lambda_\Omega \propto 1/P_{\phi,\leq 1,\Omega}$<br>**Eq 7.3**<br>$\Lambda_{B,A} = P_{\phi,\leq 1,A}/P_{\phi,\leq 1,B}$ | | | |
| Secondary outcome summarized over all four feeding attempts, namely the fraction of all mosquitoes surviving all feeding attempts required to complete their first blood meal and associated gonotrophic cycle that have been exposed to one, and only one, IRS insecticide, rather than none. | **Eq. 8**<br>$s = P_{\phi,1}\ /\ P_{\phi,\leq 1}$<br>$= P_{\phi,1}\ /\ (P_{\phi,1}+ P_{\phi,0})$<br>$= P_{\phi,1}\ /\ (P_{\phi,1}+P_{\phi,o,0,1})$ | | | |

Alternatively, the mosquito may enter a household that has been sprayed with one of the IRS insecticides used in the household-scale micro-mosaic ($n$ = 1 for $l$ = $i$ and $\alpha$ = 1), with a probability of $Q_h \pi_i$ (Figure 4 in the main manuscript). Once inside, the other sequential effects of bednet use by the human occupants come into play, the first of which is to dramatically increase the probability of being deterred ($\Delta$) from completing the feeding attempt before exposure to any active ingredients used to treat those bed nets (thereby turning them into LLINs) can occur (Figure 4 in the main manuscript). Therefore, the probability of mosquitoes being deterred and surviving over the short term but remaining hungry is the product of the baseline proportion of bloodmeals obtained from humans ($Q_h$), the baseline proportion of human bloodmeals obtained indoors ($\pi_i$) and the proportion of house-entering mosquitoes that are deterred ($\Delta$) from feeding upon the occupants inside (Table S2, Eq. 1.2).

Crucially, these are mosquitoes that will continue host seeking and commit to a second feeding attempt after being exposed to only one IRS insecticide ($n$ = 1) and could therefore reproduce if they are resistant to that single active ingredient and feed successfully in later feeding attempts without exposure to additional IRS actives ($n \geq 2$). Equation 1.2, and also those describing similarly single-exposed mosquitoes that remain "still in the game" after being deterred from feeding inside a house in later feeding attempts, are therefore highlighted with yellow shading in Table S2.

Alternatively, many mosquitoes may be taken "out of the game" because they are not deterred from attacking people indoor ($1 - \Delta$) by any NG-LLINs present and may be consequently killed by their insecticidal active ingredients ($x$ and $y$) with a much higher probability ($\mu$) than in the absence of such insecticidal nets. This lethal effect of the complementary insecticides $x$ and $y$ in the NG-LLINs reduces the number of mosquitoes that can successfully feed, survive and go on to propagate any resistance traits they may have against the single IRS active ingredient $a$ encountered in their first feeding attempt. The probability that a mosquito enters a house and dies therein as a result of attacking one or more NG-LLIN users is therefore much higher and may be calculated as the product of the baseline proportions of bloodmeals obtained from humans ($Q_h$) and doing so indoors ($\pi_i$), the proportion of mosquitoes undeterred by those bed nets ($1 - \Delta$) and the proportion of that subset consequently killed ($\mu$) immediately before or after feeding (Table S2, Eq. 1.3).

However, a subset of mosquitoes that encounter houses and enter in search of their human occupants do succeed in feeding and then survive after doing so ($f = \phi$). Although the humans in this theoretical framework are all assumed to use NG-LLINs, mosquitoes may nevertheless feed upon them indoors at various times when they are unprotected, either before they go to bed in the evening, after they get up at dawn, or during the night whenever they may or leave an exposed limb dangling outside of the net or get up briefly to attend sundry minor tasks [15, 16, 26-29]. The probability of this outcome of the first feeding attempt, resulting in a mosquito that had successfully fed and may proceed to gestate and oviposit if it is resistant to the one IRS insecticide it has encountered during its first feeding attempt, is calculated as the product of the baseline probabilities of obtaining blood from humans ($Q_h$) and doing so indoors ($\pi_i$) and the sequential probabilities of being neither deterred from feeding nor being killed in the attempt (Table S2, Eq. 1.4). Such mosquitoes that have successfully fed indoors and survived that hazardous process in their first feeding attempt are of particular interest because they have been exposed to only one IRS insecticide and may therefore pass on any resistance traits against that single active ingredient to subsequent generations. Equation 1.4, and indeed all equations describing the probabilities of mosquitoes feeding and surviving after exposure to only a single IRS active over two or more feeding attempts (Eq. 2.4.1, 3.4.1 and 4.4.1), are therefore all highlighted in dark maroon in Table S2. Note that it has always been quite normal for some small proportions of mosquitoes to feed successfully upon humans sleeping under LLINs inside houses and experimental huts, even with high quality nets and mosquito populations that are full susceptibility to the active ingredients[26, 30], so no resistance to the NG-LLIN active ingredients *x* and *y* is assumed or required to account for this small but important fraction of mosquitoes that do so.

**Possible outcomes of multiple feeding attempts necessitated by the deterrent effects of bed nets**

Of course, the probability of any particular fate ($f$) for a mosquito over multiple feeding attempts ($\alpha \geq 2$) is inherently dependent on what could have happened during its first attempt ($\alpha = 1$). Correspondingly, the calculated probabilities for the various particular intermediate or final fates detailed in Table S2 for the second, third and fourth feeding attempts ($\alpha$ = 2, 3 and 4, respectively) are all considered functions of the probabilities of deterrence away from sleeping humans, especially those protected with bednets with various properties, without obtaining a bloodmeal in the previous feeding attempt (Table S2, Eqs. 1.2, 2.2.1 plus 2.2.2 and 3.2.1 plus 3.2.2, all of which are correspondingly highlighted in yellow). In essence, the probabilities of each specific outcome listed in Table S2 for each feeding attempt are calculated as the products of each possible outcome within that feeding attempt and compound functions of the probabilities of repeatedly surviving but remaining hungry during previous feeding attempts, broken down by the number of IRS insecticides possibly encountered during foraging visits inside houses ($n$ = 1,2, 3… $N$).

As detailed in Table S2, mosquitoes completing their first feeding attempt may be exposed to no IRS insecticides if they successfully feed outdoors (Eq. 1.1) or only one IRS insecticide if they enter houses (Eqs. 1.2 to 1.4). All mosquitoes in the former category, which fed outdoors in their first feeding attempt, are clearly relevant to dampening insecticide resistance evolution because they successfully reproduce without experiencing any selection from IRS insecticides, contributing to the denominator in the secondary outcome described earlier (Eq. 7), so they are highlighted in green. However, only the minority fraction of mosquitoes that entered houses, specifically those that successfully fed and survived during their first feeding attempt, are directly and immediately relevant because they have been exposed to only one IRS insecticide (Eq. 1.4) and may propagate any resistance traits against that first-encountered active ingredient (*a*) into subsequent generations, without any lethal exposure to further IRS actives (*b*, *c*, *d*) that act as "safety net" insecticides. Those fractions of mosquitoes are consequently highlighted in maroon.

However, the variations in IRS insecticide use between different houses in a micro-mosaic (*a*, *b*, *c* etc) mean that mosquitoes are increasingly likely to be exposed to more than one IRS insecticide ($n \geq 2$) when NG-LLINs force them to undertake multiple feeding attempts ($\alpha \geq 2$). Mosquitoes deterred from blood feeding by NG-LLINs in their first attempt indoors will continue to seek hosts, often inside nearby dwellings that may be treated with a different IRS formulation (*b*, *c* etc) that kills any with resistance traits against the one they encountered on their first feeding attempt (*a*), assuming such resistance traits arise independently and randomly with negligible probability of simultaneously arising spontaneously in the same $F_0$ individual. The probability of encountering a different IRS insecticide to the first one (*b*, *c* or *d*, rather than $a$) may be expected to accumulate at a rate that is proportional to the fraction of all IRS insecticides used that these additional options account for, specifically ($N$–1)/$N$ per further feeding attempt, assuming equal and random coverage with each of these different IRS options. Thus, the risks of cumulative exposure to two or more IRS insecticides ($n \geq 2$) over multiple feeding attempts ($\alpha \geq 2$) are accounted for in equations 2.2.2, 3.2.2 and 4.2.2 (Mosquitoes deterred from feeding on their second, third and fourth attempts, respectively), as well as equations 2.4.2, 3.4.2 and 4.4.2 (Mosquitoes that successfully fed and survived their second, third and fourth attempts, respectively), all of which are consequently highlighted in blue. On the other hand, the increasingly lower probabilities of repeated exposure to the same IRS insecticide over multiple visits to houses are calculated as compound functions of 1/$N$ in Eqs. 2.2.1, 3.2.1 and 4.2.1, plus 2.4.1, 3.4.1 and 4.4.1, all of which are highlighted in maroon as a sign of potential danger.

**Numerical validation**

Equations 1.5, 2.5, 3.5, and 4.5 represent logical relationships between all the different possible mosquito fates arising from each possible feeding attempt, allowing numerical validation that all possible fates are fully accounted for and add up to logically sensible exact totals. The inflow of mosquitoes into a given feeding attempt equals the total outflow across all their subsequent potential outcomes, successful feeding indoor or outdoor, deterrence or mortality.

Equation 1.5 applies to the very first feeding attempt, wherein the entire mosquito population starts foraging for blood with an assumed probability of exactly one. At this stage, every mosquito must progress to one of four mutually exclusive fates that are separately calculated, namely (1) successfully feeding outdoors with no IRS exposure ($f = \phi$, $l = o$, $n = 0$), (2) entering a house and being deterred by any bednets present but only after also gaining one exposure to a single slow-acting IRS insecticide ($f = \Delta$, $l = i$, $n = 1$), (3) being killed indoors by the bednets before that single exposure to a slow-acting IRS insecticide can take effect ($f = \mu$, $l = i$, $n = 1$), or (4) successfully feeding indoors, again after gaining one IRS exposure to a single insecticide within that one house ($f = \phi$, $l = i$, $n = 1$). Since these outcomes cover all the possibilities within the conceptual framework of the model (Figure 4 in the main manuscript) for the first feeding attempt without any overlap, their probabilities must sum to exactly 1. Equation 1.5 was therefore used to confirm numerically that no mosquito fates were miscalculated, lost track of or double counted in the model predictions for the first feeding attempt.

Equation 2.5 totals all the possible outcomes of mosquitoes making a second feeding attempt, assuming that these possibilities can apply only to the mosquitoes that were deterred from feeding in the first attempt ($P_{\Delta,i,1,1}$) and must so make such a second attempt. These mosquitoes now face the same set of possible fates, but their outcomes are split by whether they accumulate a second IRS insecticide that is distinct from the first ($n = 2$, after exposure to both $a$ and $b$ in the first and second of two houses visited) or remain with exposure to only such slow-acting IRS insecticide ($n = 1$, after either only visit to a single house containing IRS insecticide $a$ or two visits to two houses that both contained $a$). Equation 2.5 therefore breaks the proportion of mosquitoes deterred in the first feeding attempt down into similar terms to those used for all mosquitoes in equation 1.5, except that some of these categories are further broken down according to how many slow-acting IRS insecticides those mosquitoes were exposed to while visiting houses: (1) Outdoor feeding, (2) deterrence (but now split according to whether $n = 1$ or 2), (3) indoor mortality, or (4) indoor feeding (now also split by $n = 1$ or 2).

This exact equality ensures complete accounting for the probabilities for various subsets of mosquitoes continuing to the second attempt by simply summing the calculated values for each of them and comparing with the logical expectation outlined by equation 2.5.

For equation 3.5, it follows that the mosquitoes deterred from the second attempt will necessarily make a third one, at the end of which they may have experienced cumulative exposure to as many as three slow-acting IRS insecticides ($n$ = 1, 2 or 3 in scenarios where $N$ = 3, specifically $a$, $b$ and $c$). The fates are again broken down exhaustively to account for all mosquitoes entering the third feeding cycle, this time broken down by diversity of IRS insecticide exposure for all outcomes except immediate indoor mortality: (1) Outdoor feeding (but now split by $n$ = 1 or ≥2), (2) deterrence from feeding yet again (again, split by $n$ = 1 or ≥2), (3) indoor mortality regardless of IRS exposure history, and (4) indoor feeding (again, split by $n$ = 1 or ≥2). The more complex tracking of IRS insecticide exposure diversity as a compound function of the diversity of the micro-mosaic itself ($N$) introduces far greater opportunities for mathematical and computational error, so it becomes even more important to ensure that the sum of all third-attempt outcome probabilities must exactly match the inflow from those diverted in the second attempt, verifying that the model correctly handles accumulating insecticide diversity without loss or gain.

Lastly, equation 4.5, covers the fourth feeding attempt, and the last considered in this model implementation for practical reasons of parsimony, relating only to the quite small fractions of mosquitoes expected to be deterred from their third feeding attempts. The same exhaustive breakdown by clearly distinct fates in 3.5 are applied one last time, stratified by history of exposure to different slow-acting IRS insecticides (n=1 or ≥2).

Applying equations 1.5, 2.5, 3.5 and 4.5 to all the various outcomes that were considered possible based on the assumed conceptual framework (Figure 4 in the main manuscript), all of the equations detailed in the rows of Table S2 immediately above them, as well as the spreadsheet modelling tools used to apply them in practice were fully validated: All calculated values were confirmed to matched exactly to those expected.

**Summary primary, secondary and explanatory outcomes**

Equation 6 calculates the primary outcome, which is the total proportion of all mosquitoes that ultimately succeed in obtaining a blood meal after having been exposed to no more than one IRS insecticide through one or more feeding attempts in their first gonotrophic cycle ($P_{\phi,\leq 1}$). This is achieved by summing eight specific probability terms that represent all possible pathways leading to successful feeding that result in exposure to only one IRS insecticide or none at all, making $P_{\phi,\leq 1}$ directly proportional to the rate at which $F_0$ mosquitoes with new incipient resistant traits may be expected to survive exposure to that active ingredient and propagate those resistance traits to subsequent $F_{\geq 1}$ generations, despite any effects of IRS micro-mosaics and/or NG-LLINs.

In terms of our wildfire analogy, this primary outcome may be conceptualized as the mosquitoes that feed outdoors without exposure represent sparks that land beyond the reach of suppression by indoor interventions, while those exposed to exactly one IRS insecticide represent sparks that encounter a single control barrier yet persist. Together, these pathways define the conditions under which the first flames of resistance and onward transmission may continue to flicker, within landscapes managed by IRS micro-mosaics and NG-LLIN interventions.

Assuming that the combinations of IRS and NG-LLINs deployed together render each other redundant in terms of vector population suppression prior to the emergence of novel resistant traits, and that full coverage is necessarily sustained to consistently maintain the minimum vector population size possible, then the rate at which such incipient traits emerge ($\varepsilon$) from $F_0$ to $F_{\geq 1}$ in a given intervention scenario ($\Omega$) is proportional to the absolute survival and reproduction probability under those conditions ($P_{\phi,\leq 1,\Omega}$), as per equation 7.1). By corollary, the expected lifetime of pre-emptive use for that insecticide, before a resistance trait against it emerges from $F_0$ to $F_{\geq 1}$ and can subsequently be selected for, is inversely proportional to the absolute survival and reproduction probability in that intervention scenario ($\lambda_\Omega$), as per equation 7.2. Correspondingly, the relative duration of the expected mean lifetime for pre-emptive use for an insecticide ($\Lambda_{B,A}$) in one pre-emptive intervention scenario versus another ($\Omega$ = B relative to A, as per the third versus second panel of Figure 3 in the main manuscript) may be calculated as the reciprocal of the quotient of their absolute survival and reproduction probabilities, as per equation 7.3.

For the secondary outcome, equation 8 calculates the widely accepted evolutionary parameter defined as the selection coefficient ($s$) [20, 21], which is the fraction of all mosquitoes that successfully obtain a blood meal and will survive to complete their gonotrophic cycle that have been exposed to exactly one IRS insecticide that they are resistant to, rather than to none. The denominator of equation 8 thus represents the total proportion of the population that will feed successfully assuming they are resistant to a single IRS insecticide, and the overall selection coefficient term reflects the evolutionary pressure acting upon such traits that drive growing frequency trends, possibly culminating in fixation. In terms of our wildfire analogy, this secondary outcome may be conceptualized as the intensity with which the wildfires burn once they have been initiated by sparks of incipient traits that various intervention combinations may fail to suppress at source.

Beyond all the different fates predicted for each feeding attempt in the first part of Table S2, these various outcomes were also categorized by *final* fate ($f = \phi$ and $\mu$ but not $\Delta$), location ($l$ = $i$ or $o$) and IRS insecticide exposure history ($n$ = 0, 1 or ≥2) and summed across all feeding attempts to get overall summaries of how these probabilities accumulated over entire gonotrophic cycle histories. These cumulative probabilities of final fates allow the overall influences of bednet interventions that may sometimes only defer preferred and non-preferred outcomes until after two or more feeding attempts to be understood more synthetically than from the breakdowns by individual sequential feeding attempts.

**Model parameterization to represent broadly relevant stereotypes of vector behaviour and bednet properties**

Parameterization of this mosquito feeding model requires assigning empirically validated values to key behavioural parameters derived from field observations in deliberately approximate, readily grasped terms that therefore represent stereotypes of two broad categories of vectors from across Africa. For *An. funestus*, which was chosen as a particularly stereotypical representative of a highly anthropophagic and endophagic malaria vector, in which feeding behaviour is strongly skewed toward humans and occurs predominantly indoors during late night hours when most people are asleep and can protect themselves with a bed net [24, 31, 32]. Evidence from multiple sub-Saharan African settings consistently demonstrates a high human blood index, even in areas with widespread deployment of indoor interventions such as LLINs and IRS [2, 5], underscoring the species' persistent preference for human hosts [24, 31, 32]. Correspondingly, a 90% probability was assumed for this species encountering and try to attack a human rather than an animal in any given feeding attempt ($Q_h$ = 0.90). Similarly, a 90% probability was assumed for human-directed attacks occurring indoors in the absence of bednets ($\pi_i$= 0.90) [2, 5, 16], reflecting strong endophagic tendencies.

In contrast to *An. funestus*, *An. arabiensis* exhibits pronounced behavioural plasticity, flexibly exhibiting both anthropophagy and zoophagy, as well as endophagy and exophagy with approximately similar preference [24, 31, 32]. This species frequently seeks blood meals from cattle in particular and often bites outdoors during early evening or late morning, substantially reducing its contact with indoor interventions such as LLINs and IRS [24, 31, 32]. Similarly to other models of malaria transmission dynamics, here we assume an intermediate human blood index of 50% ($Q_h$ = 0.5) in the presence of LLINs/IRS [13, 23, 33, 34] for the stereotypical *An. arabiensis* modelled herein, reflecting its comparable preferences for human and non-human hosts. Additionally, only half of the human-directed feeding attempts occurring the absence of bednets were assumed occur indoors, as shown by a 50% baseline probability that an attack upon a human will occur indoors ($\pi_i$= 0.5) [1, 17, 35].

*An. funestus* is vulnerable to indoor interventions such as LLINs and IRS, exhibiting relatively little caution when it attempts to feed on humans indoors or rest indoors before and/or after feeding [31, 32]. Field evaluations of LLINs consistently report high mosquito mortality, so an immediate indoor mortality rate of 80% was assumed for *An. funestus* attempting to feed inside houses ($\mu$ = 0.80) across all NG-LLIN scenarios, similar to those measured in several experimental hut trials [25, 36-38] . For NG-LLINs nets incorporating lethal insecticidal actives only, 60% deterrence was assumed for this species ($\Delta$ = 0.60) based on available field estimates for untreated bednets that provide only physical protection [36]. In contrast, NG-LLINs combining excito-repellent and lethal actives were assumed to exhibit substantially higher deterrence ($\Delta$ = 0.80), consistent with field measurements for typical pyrethroid-based LLINs [25, 36]. In the absence of any bed net intervention, mosquitoes were nevertheless assumed to experience low baseline levels of 10% for both natural mortality and deterrence within houses ($\Delta$ and $\mu$ = 0.10), as in previous models similarly parameterized base on overviews of existing field evidence [17, 39].

However, even when the more zoophagic and exophagic species *An. arabiensis* does enter houses, it tends to do so more cautiously than *An. funestus* and leave again more promptly when it fails to obtain a bloodmeal therein [23, 40]. Field-based assessment of LLINs indicate somewhat higher rates of exiting without feeding, and correspondingly more moderate mortality rates for *An. arabiensis* inside experimental huts than for more anthropophagic species like *An. gambiae* and *An. funestus*, presumably reflecting its lower contact rates with treated surfaces. In keeping with the results of those field studies [23, 40], we therefore assumed slightly increased deterrence rates of 70% for untreated bednets and NG-LLINs with only lethal insecticides and 90% for NG-LLINs that include insecticides with excito-repellent properties ($\Delta$ = 0.7 and 0.9, respectively) were assumed for this more evasive vector [17, 36], which is considered a stereotypical representative of numerous vectors around the tropics that dominate residual malaria transmission after LLINs and/or IRS have been scaled up [3].
